# Engineering orthogonal quorum sensing circuits using LuxR-type systems in yeast consortia

**DOI:** 10.64898/2026.05.13.724834

**Authors:** Aafke C.A. van Aalst, Maxence Holtz, Michal Poborsky, Christoph Crocoll, Emil Damgaard Jensen, Michael Krogh Jensen

**Author notes:** correspondence: Aafke C.A. van Aalst / Present address: Department of Process and Life Science Engineering, Lund Tekniske Högskola (LTH), Naturvetarvägen 22, 22100 Lund, Sweden, Emil Damgaard Jensen.

## Abstract

Engineered microbial communities hold significant biotechnological potential because their collective metabolism can produce functions beyond those achievable by individual strains. However, multicellular synthetic gene circuits require orthogonal communication systems that enable precise, programmable signaling between cells. Quorum sensing (QS), where cells both produce and detect small diffusible signal molecules, offers a natural framework for such intercellular communication. However, the construction of complex multicellular circuits for applications such as biobased production is currently hampered by the limited number of orthogonal QS channels available in yeast. Here, we expand the QS toolkit in *Saccharomyces cerevisiae* by characterizing four LuxR-type biosensors based on EsaR, LasR, TraR and RpaR, alongside the previously established LuxR biosensor. We functionally expressed acyl-CoA-dependent HSL synthases in yeast, producing a diverse range of aliphatic and aromatic HSL signals. LuxR and RpaR, were compatible with *in vivo* ligand production and established as orthogonal QS signaling pair with synthases MesI and RpaI, respectively. Co-culture experiments demonstrated QS-dependent intercellular signaling, with 3.9-fold and 6.4-fold induction relative to monocultures. Together, these results establish a modular and extensible platform for orthogonal intercellular communication in yeast, enabling the construction of multicellular synthetic gene circuits.

## Introduction

Inspired by microbial natural consortia, rationally designed microbial synthetic consortia have extended from food fermentation processes to emerge as integral solutions to improve bioproduction. Such systems allow for better co-factor and intermediates balancing ^1-3^, reduce pathway metabolic imbalances through division of labour ^2-5^ and provide optimal environments best suited for different pathway enzymes ^6,7^. To coordinate behaviour of individual subpopulations within these consortia, reliable tools for communication between microbial cells are required. These tools enable populations to exchange information, implement feedback and stabilize community composition ^8, 9^.

In nature, population-wide communication is often achieved through quorum sensing (QS). QS is a process in which cells continuously transmit and detect signaling molecules in order to control specific cellular processes based on the concentration of other cells in the environment. Acylated homoserine lactones (AHLs) are small signaling molecules used by many Gram-negative bacteria for cell-to-cell communication ^10-12^. Most AHLs consist of a homoserine lactone (HSL) ring as a core and a fatty acid-derived acyl side chain. These fatty acyl-HSLs are mostly synthesized by synthases belonging to the LuxI protein family. Because of the different sources of the acyl side chain, the molecule can vary in length, oxidation state and functional groups ^13, 14^. The structural variation of homoserine lactones leads to a large natural diversity in both signaling molecules and corresponding regulators. For example, the signaling molecule 3-oxo-C12-HSL is produced by LasI and detected by LasR, while 3-oxo-C6-HSL is native to the LuxI-LuxR system. Several of these LuxI-LuxR-type systems have been employed in synthetic biology applications ^15, 16^, including for microbial consortia engineering ^17, 18^. Several natural QS pairs have been combined and characterized to generate signal-orthogonal QS systems in bacteria, in which a receptor responds exclusively to its cognate AHL. Examples include LuxI/LuxR and LasI/LasR ^19^, EsaI/TraR and LasI/LasR ^15^, LuxI/LuxR and RpaI-RpaR ^20^, BjaI/BjaR and EsaI/TraR ^16^ and EsaI/EsaR and LasI/LasR ^21^.Signal orthogonality enables multiple communication channels to operate simultaneously within a single environment without unwanted cross-activation.

In yeast-based consortia, however, population coordination has largely relied on metabolic interactions such as cross-feeding ^22-24^, with only a few engineered intercellular communication systems ^25, 26^. Moreover, the implementation of AHL-based quorum-sensing components in yeast remains limited: to date, only LuxR has been functionally reconstituted in *Saccharomyces cerevisiae* ^27, 28^ and only a single AHL, i.e.C8-HSL, has been produced, and at low titers ^27^. This limitation is partly attributed to differences in fatty acid metabolism between yeast and bacteria, as acyl carrier protein (ACP) in yeast is tightly associated with fatty acid synthase complexes ^29-31^, potentially restricting the activity of ACP-dependent LuxI enzymes. In this study, we aim to increase the number of functionalized AHL-based QS-systems in yeast to obtain signal-orthogonal QS pairs. Development of an AHL-based orthogonal intercellular communication system in yeast would be directly applicable in yeast-yeast or yeast-bacteria co-cultures as well as yeast-bacteria.

To establish novel QS systems, we introduce and optimize several AHL-biosensors as well as CoA-dependent synthases. Additionally, production of phenyl-acetyl-HSL is engineered through introduction of biosynthetic enzymes, and enzyme localization strategies are employed to optimize AHL biosynthesis. Together, this work expands the repertoire of AHL signals that can be produced and sensed in yeast, providing a foundation for reliable engineering of communication in synthetic microbial consortia.

## Results and discussion

### Engineering repressor-type EsaR-based biosensors

For the development of AHL-biosensors we first investigated the transcriptional repressor EsaR, which has been described to respond to 3-oxo-C6-HSL ^32, 33^, incombination with its canonical operator sequence (*esaO*; GCCTGTACTATAGTGCAGGT) which we inserted downstream of the TATA-box in p*TEF1* (strains AAA015; p*TEF1*_*esaO*-yeGFP and AAA033; p*TEF1*_*esaO*-yeGFP, p*PGK1*-*esaR*). When comparing yeGFP fluorescence of strains AAA015 and AAA033, no repression of yeGFP signal was observed in the presence of repressor EsaR (Figure S1). This indicates that EsaR is not fully functional in yeast.

Previous studies in *E. coli* identified EsaR variants V220A and D91G ^34^ as mutations that increase sensitivity and we tested whether these mutations in yeast could improve activity of EsaR (strains AAA031; p*TEF1*_*esaO*-yeGFP, p*PGK1*-*esaR*_V220A_ and AAA032; p*TEF1*_*esaO*-yeGFP, p*PGK1*-*esaR*_D91G_). Both mutant repressors were able to repress yeGFP expression in the absence of ligands, by an approximate 2.6- and 2.7-fold repression respectively (Figure S1). However, only the D91G variant enabled de-repression following addition of AHL-compounds (100 µM) (Figure S1). The V220A mutation is located in the DNA-binding domain ^34^, which may contribute to enhanced repression, though whether ligand binding and allostery are preserved remains unclear. D91G on the other hand is located in the AHL-binding domain ^34^, which may play a role in maintaining or restoring ligand responsiveness in yeast.

EsaR_D91G_-based biosensor responded to 3-oxo-C6-HSL, 3-oxo-C8-HSL, 3-oxo-C12-HSL, C6-HSL and C8-HSL (Figure S1). When tested specifically with 3-oxo-C6-HSL, a 1.8-fold increase in yeGFP fluorescence was observed at 100 µM (Figure 1A). To further increase the dynamic range of the biosensor, we next targeted the hybrid promoter architecture. We inserted an additional copy of *esaO* downstream of the TATA-box in p*TEF1* (strains AAA145; p*TEF1*_2x*esaO*-yeGFP and AAA167; p*TEF1*_2x*esaO*-yeGFP, p*PGK1*-*esaR*_D91G_).

**Figure 1:**
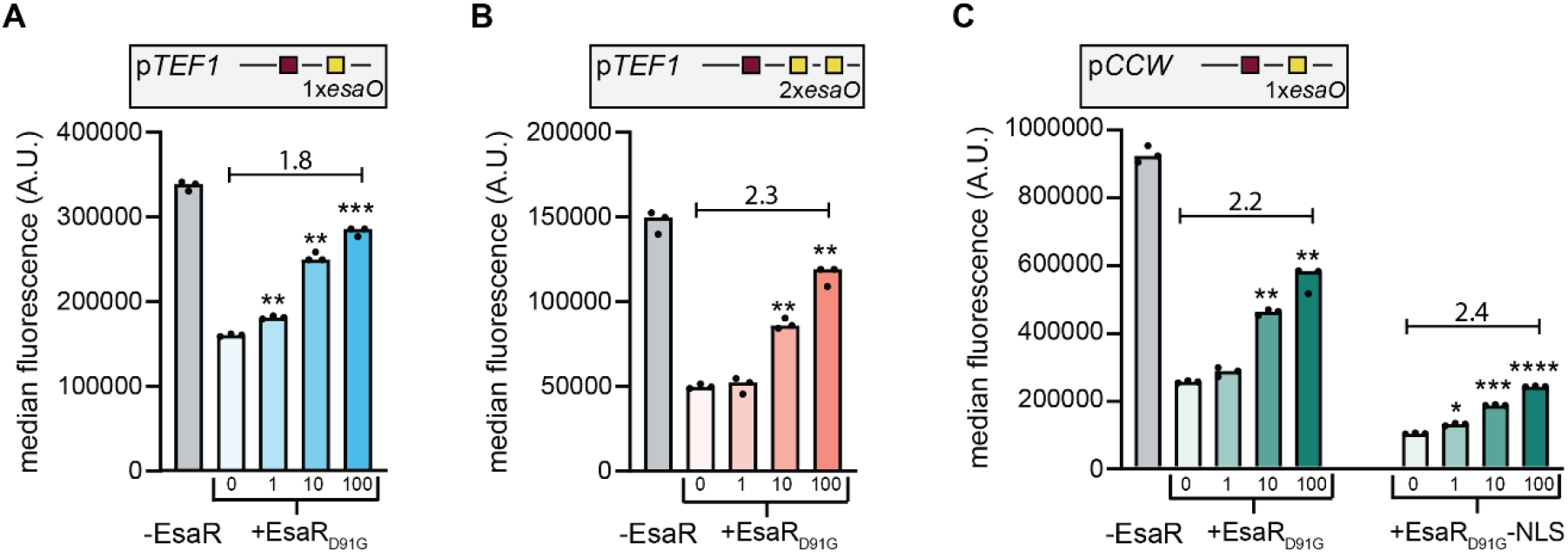
Engineering of repressor-type AHL biosensors. yeGFP median fluorescence determined by flow cytometer 6 h after supplementation with 0, 1, 10 or 100 µM of 3-oxo-C6-HSL. Number above the line shows maximum dynamic range expressed as fold change between 0 and 100 µM ligand as indicated. **(A)** -EsaR indicates strain AAA015 (p*TEF1*_*esaO1*-yeGFP) and +EsaR_D91G_ indicates strain AAA032 (p*TEF1*_*esaO1*-yeGFP, p*PGK1*-*esaR*_D91G_). **(B)** –EsaR indicates strain AAA145 (p*TEF1*_2x*esaO1*-yeGFP) and +EsaR_D91G_ indicates strain AAA167 (p*TEF1*_2x*esaO1*-yeGFP, p*PGK1*-*esaR*_D91G_). **(C)** -EsaR indicates strain AAA213 (p*CCW*-*esaO*-yeGFP), +EsaR_D91G_ strain AAA216 (p*CCW*-*esaO*-yeGFP, p*PGK1*-*esaR*_D91G_) and +EsaR_D91G_-NLS strain AAA218 (p*CCW*-*esaO*-yeGFP, p*PGK1*-*esaR*_D91G_-NLS). Promoter representations: red box indicates TATA-box, operator sequence(s) depicted as yellow box(es) (*esaO*). A.U.: arbitrary units. Individual points of biological replicates (n=3) are shown. Significance of difference between induced (1, 10 or 100 µM) and uninduced (0 µM) is indicated above each bar as follows: ^*^ : p-value ≤ 0.05, ^**^ : p-value ≤ 0.01, ^***^ : p-value ≤ 0.001 and ^****^ : p-value ≤ 0.0001.

This resulted in 2.3-fold lower basal activity of the promoter and a 3.4-fold reduction in yeGFP expression in the presence of the regulator and absence of 3-oxo-C6-HSL. Addition of 100 µM ligand led to a ∼70% derepression and a 2.3-fold induction of yeGFP expression (Figure 1B). To test if we could further improve the dynamic range, we subsequently inserted a single *esaO* copy into p*CCW12* (hereafter p*CCW*) as core promoter (strains AAA213; p*CCW*-*esaO*-yeGFP and AAA216; p*CCW*-*esaO*-yeGFP,p*PGK1*-*esaR*_D91G_). Basal yeGFP expression from p*CCW* was 2.5 and 5.7-fold higher than that observed for p*TEF1*-derived promoters. Regulator expression reduced reporter output by 3.8-fold, and ligand addition mediated ∼60% of derepression (Figure 1C).

Alternatively, we tested the addition of a nuclear localization signal, using the p*CCW*-based system (strain AAA218: p*CCW*-*esaO*-yeGFP, p*PGK1*-*esaR*_D91G_-NLS). This further enhanced repression in the absence of ligand up to 9.8-fold. This can most likely be explained by the fact that more repressors reach the operator sequence (Li 2015).However, adding the ligand only resulted in a 25% derepression and a 2.4-fold induction of yeGFP (Figure 1C).

While these different configurations all achieved similar dynamic ranges (1.8–2.4-fold), they differed substantially in absolute yeGFP expression levels. The two p*TEF1*-derived designs spanned ∼160,000–2,800,000 A.U. and ∼50,000–120,000 A.U., while the two p*CCW*-derived designs spanned ∼250,000–580,000 A.U. and ∼100,000–240,000 A.U., respectively. This highlights the tunability of promoter–repressor architectures for application-specific requirements (Figure 1). The observed variation in absolute expression levels despite similar dynamic ranges highlights the tunability of promoter– regulator architectures. This flexibility allows biosensor designs to be tailored to application-specific requirements, such as minimizing background signal or maximizing output intensity.

### Development of AHL-biosensors using transcriptional activators

Previously, we engineered a highly sensitive AHL-biosensor based on the transcriptional activator LuxR, with dynamic ranges exceeding 100-fold ^27^. We therefore decided to test out other QS transcriptional activators TraR, LasR and RpaR. In order to attempt to boost transcriptional activation, each activator was fused to the VP48 activation domain and expressed together with a synthetic promoter containing five operator sites driving the expression of yeGFP (*traO* ^35^; ATGTGCAGATCTGCACAT, *lasO* ^36^;ATCTATCTCATTTGCTAGTT, *rpaO* ^37^; ACCTGTCCGATCGGACAGTA). For clarity, these are hereafter referred to as TraR-, LasR- and RpaR-based biosensors respectively.The LasR-based biosensor (AAA269; 5x*lasO*-yeGFP, AAA274; 5x*lasO*-yeGFP, p*PGK1*-VP48-*lasR*) responded robustly to both C12-HSL and 3-oxo-C12-HSL, exhibiting a strong yeGFP induction of up to 358-fold (Figure 2A). Similarly, the RpaR-based biosensor (AAA160; 5x*rpaO*-yeGFP, AAA258; 5x*rpaO*-yeGFP, p*PGK1*-VP48-*rpaR*) showed specific activation in response to its canonical ligand, *p*-coumaroyl-HSL of up to 34-fold (Figure 2B). Lastly, TraR (AAA271; 5x*traO*-yeGFP, AAA75; 5x*traO*-yeGFP, p*PGK1*-VP48-*traR*) displayed responsiveness towards its canonical ligand 3-oxo-C8-HSL, though with a relatively lower maximum yeGFP induction of up to 20–fold (Figure 2C).

**Figure 2:**
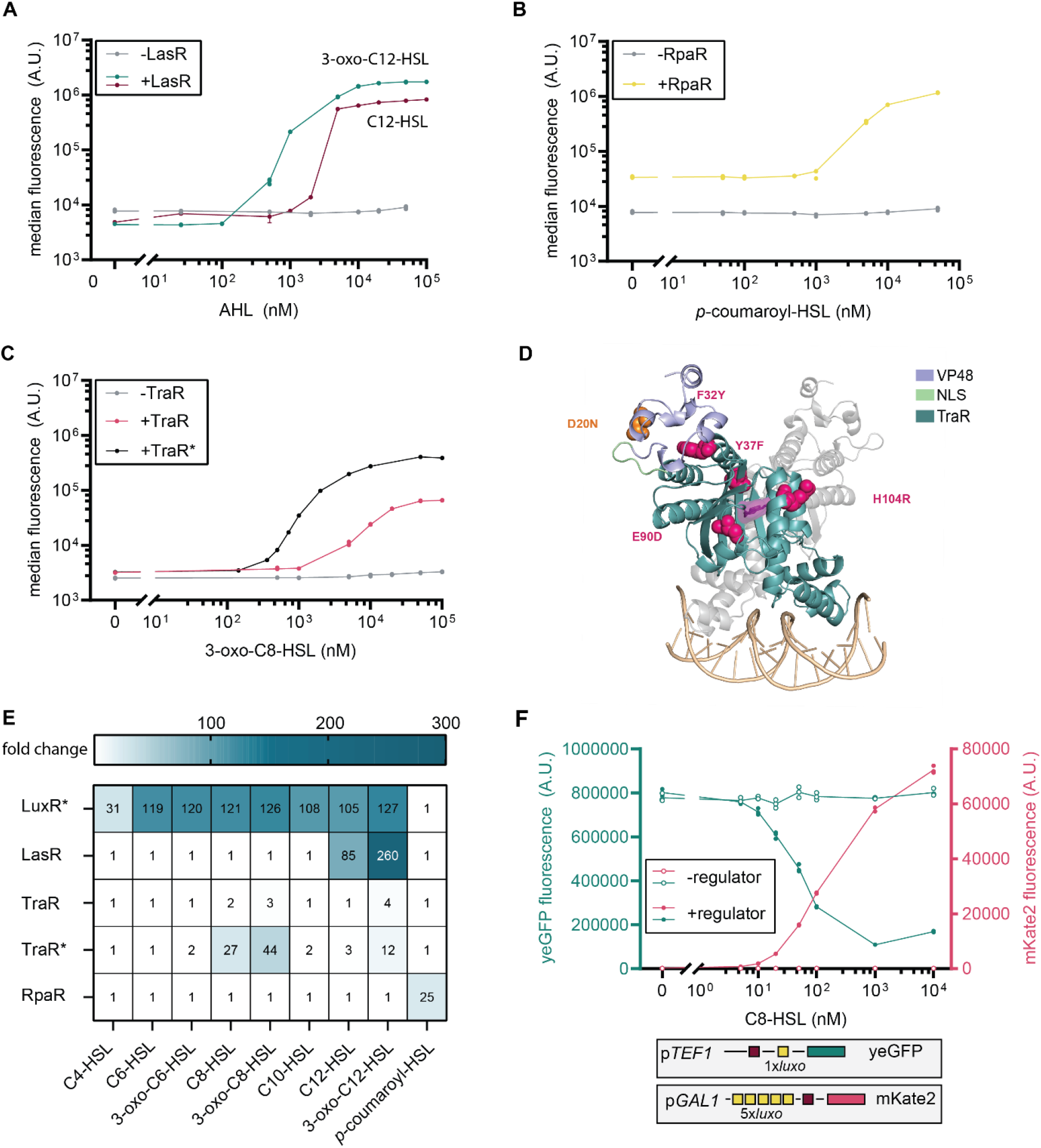
Engineering QS-biosensors using activator-type regulators. **(A)** Dose-response curves of yeGFP fluorescence levels 6 h following supplementation of 0–100,000 nM 3-oxo-C12-HSL and C12-HSL, tested for strains AAA269 (-LasR: 5x*lasO*-yeGFP) and AAA274 (+LasR: 5x*lasO*-yeGFP, p*PGK1*-VP48-*lasR*). **(B)** Dose-response curves of yeGFP fluorescence levels 6 h following supplementation of 0–100,000 nM *p*-coumaroyl-HSL, tested for strains AAA160 (-RpaR: 5x*rpaO*-yeGFP) and AAA258 (+RpaR: 5x*rpaO*-yeGFP, p*PGK1*-VP48-*rpaR*). **(C)** Dose-response curves of yeGFP fluorescence levels 6 h following supplementation of 0–100,000 nM 3-oxo-C8-HSL, tested for strains AAA271 (-TraR: 5x*traO*-yeGFP), AAA275 (+TraR: 5x*traO*-yeGFP, p*PGK1*-VP48-*traR*) and ACA080 (+TraR*: 5x*traO*-*amdS*, 5x*traO*-yeGFP, p*PGK1*-VP48-*traR\**). **(D)** Structural mapping of mutations identified in TraR*. The structural model was generated using Boltz-2 and includes a TraR* dimer co-folded with the ligand 3-oxo-C8-HSL (purple) and the DNA operator *traO* (yellow). **(E)** Heatmap indicating the fold change of yeGFP induction measured by flow cytometer 6 h after supplementation with 0 and 5 µM of ligand of biological duplicates. LuxR* indicates strain AAA156 (5x*luxO*-yeGFP, p*PGK1*-*GAL4*_AD_-*luxR*\*)^27^. **(F)** Dose-response curves of yeGFP and mKate2 fluorescence levels 6 h following supplementation of 0–10,000 nM C8-HSL, measured for strains AAA295 (5x*luxO*-mKate2, p*TEF1*_*luxO*-yeGFP) and AAA289 (5x*luxO*-mKate2, p*TEF1*_*luxO*-yeGFP, p*PGK1*-*GAL4*_AD_-*luxR\**). Promoter representations: red box indicates TATA-box, operator sequence(s) depicted as yellow box(es) (*luxO*). For A, B, C, F: Individual points of biological replicates (n=3) are shown. A.U.: arbitrary units.

### Improving dynamic range and sensitivity of TraR

To improve the performance of the TraR-based biosensor, we screened a TraR mutant library using previously established growth-based selection pipeline developed in our laboratory to enrich variants with increased dynamic range ^27^. Selection was performed at 1 µM 3-oxo-C8-HSL, a concentration just below the operational range of the parental biosensor, thereby simultaneously enriching for increased sensitivity. Growth-based selection was implemented using the *amdS* marker, while population-level enrichment was monitored via yeGFP fluorescence. Following selection in the presence of 1 µM 3-oxo-C8-HSL, a more fluorescent population was observed (Figure S2) and subsequently plated to isolate individual variants. Six colonies were analyzed, all of which had an identical genotype (TraR_mut5, subsequently referred to as TraR*). The performance of the enriched TraR* variant (ACA080) was evaluated by dose–response analysis using 3-oxo-C8-HSL (Figure 2C). Compared to the parental biosensor, the mutant exhibited both an increased dynamic range and enhanced sensitivity. At 100 µM, the yeGFP induction was increased from 20 to 68-fold, while significant sensor signal at 0.5 µM was also observed.

We identified 5 mutations, of which one in the activation domain (Table 1, Figure 2D). E90D and H104R are located within the region assigned for the AHL-binding domain (residues 39-140) ^38^. Structural models generated with Boltz2 ^39^ and co-folded with the 3-oxo-C8-HSL ligand indicate that these mutations do not appear to affect ligand binding (Figure S3A). The binding pockets conformation and ligand-protein interactions with key residues identified in the available TraR crystal structure (PDB: 1H0M) remain unchanged (Figure S3B). No mutations were found at the dimerization interface,indicating it is likely unaffected. The F32Y mutation introduces a new hydrogen bond with the His3 residue in a neighboring helix, which may influence allostery (Figures S3C).We could not find obvious explanations for the other mutations, which may affect protein expression, dynamics, folding, or be neutral passengers from the error-prone PCR without clear functional impact.

**Table 1:** overview of mutations found in VP48-*traR* in selected mutant.

| Location | Mutations |
| --- | --- |
| VP48 | D20N (GAC→AAC) |
| TraR | F32Y (TTC→TAC), Y37F (TAT→TTT), E90D (GAA→GAT), H104R (CAT→CGT) |

**Table 2:**
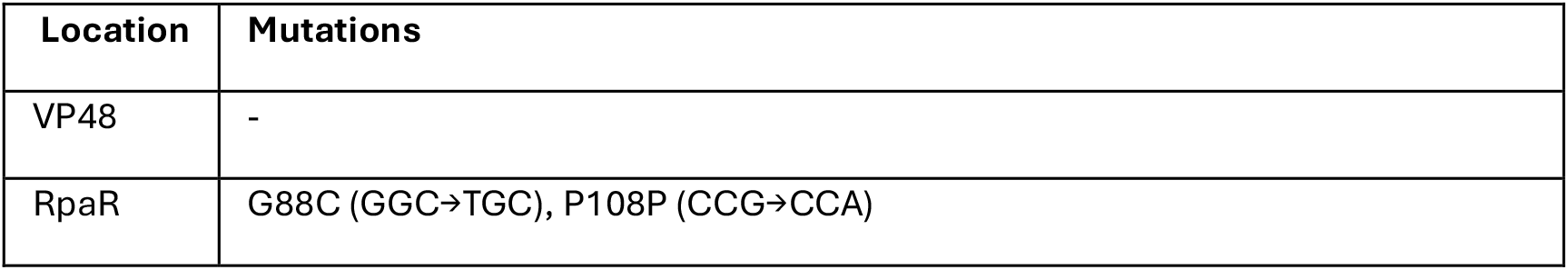
overview of mutations found in VP48-*rpaR* in selected mutant.

### Testing the specificity of the biosensors

For orthogonal biosensor systems, understanding cross-reactivity and ligand specificity is essential. We therefore evaluated the response of our newly developed activator-based biosensors to a panel of AHL compounds at 5 µM and included the previously developed LuxR-based biosensor for comparison ^27^. Previously, three LuxR-mutants were selected, each carrying the N86 mutation. We used the mutant previously referred to as Gal4_AD-LuxR_gen1 ^27^, which we will refer to as LuxR* throughout this study. LuxR* responded to all tested AHLs except *p*-coumaroyl-HSL, consistent with its broad ligand promiscuity (Figure 2E) ^40^. In contrast, LasR and TraR* displayed higher specificity, responding primarily to their canonical ligands (3-oxo-C12-HSL and 3-oxo-C8-HSL, respectively) as well as to their corresponding non-oxygenated variants (C12-HSl and C8-HSL), albeit with reduced yeGFP induction (Figure 2E). TraR* additionally showed partial responsiveness to 3-oxo-C12-HSL. RpaR exhibited strict specificity and was exclusively activated by *p*-coumaroyl-HSL (Figure 2E). Together, these ligand specificity profiles define the orthogonality potential of the four biosensors. Since RpaR responds exclusively to aryl-HSL signals, it can be paired orthogonally with any of the three AHL-responsive biosensors. Additionally, despite partial cross-reactivity of TraR* towards 3-oxo-C12-HSL, LasR and TraR* recognize sufficiently distinct ligands to form an orthogonal pair.

### Dual-mode transcriptional system using Gal4 activation domain

In its native context, LuxR most often functions as a transcriptional activator by recruiting RNA polymerase to the promoter, though it can also repress gene expression by preventing RNA polymerase binding or by impeding its movement along the promoter, depending on where the operator sequence is located ^41, 42^. In our previous study we demonstrated that transcriptional output direction of our LuxR-based biosensor could be controlled by inclusion or omission of the VP48 activation domain, necessitating two distinct regulator configurations for activation and repression ^27^. While our biosensors based on TraR*, LasR and RpaR are fused with VP48 activation domain, LuxR* is fused to the less potent Gal4-activation domain. To enable bimodal transcriptional control in which a single regulator configuration could both activate and repress transcription depending on promoter architecture within one cell, we evaluated the use of Gal4 activation domain instead. We combined LuxR* (Gal4_AD-LuxR_gen1) with a repression-based promoter driving yeGFP (p*TEF1*_*luxO*-yeGFP) expression and an activation-based promoter driving mKate2 expression (p*GAL*_core_5x*luxO*-mKate2) (AAA295; p*TEF1*_*luxO*-yeGFP, p*GAL*_core_5x*luxO*-mKate2 and AAA289; p*TEF1*_*luxO*-yeGFP, p*GAL*_core_5x*luxO*-mKate2, p*PGK1*-*GAL4*_AD_-*luxR*_gen1). Fluorescence output was measured following supplementation with increasing concentrations of C8-HSL (Figure 2F). In contrast to VP48, using a less active activation domain enabled both ligand-dependent repression and activation mediated by the same transcriptional regulator within a single cell.

### Introduction of acyl-CoA-dependent synthases in Saccharomyces cerevisiae

After establishing new AHL-biosensors in yeast, with different specificities, we next decided to expand the repertoire of AHL production to characterize biosensing based on endogenous ligand formation. We focused on introducing synthases that have been described to be CoA-dependent, since yeast does not have freely available acyl-ACP to donate an acyl-group ^29-31^. Based on substrate preference and structural features, CoA-dependent AHL synthases have been divided into 4 clusters ^43^ (Figure 3A) and we selected synthases from the different clusters and ordered these sequences codon-optimized. From cluster 1a, we chose BjaI from *Bradyrhizobium japonicum*, which exclusively uses isovaleryl-CoA instead of isovaleryl-ACP ^13, 44, 45^ (Figure 2A). We picked two enzymes from cluster 1b; BraI from *Bradyrhizobium* which has been described to be specific for cinnamoyl-CoA ^46^, and RpaI from *Rhodopseudomonas palustris* which uses *p*-coumaroyl-CoA ^37^ (Figure 3A). From cluster 1c, MesI was selected, which shows the highest catalytic efficiency with hexanoyl-CoA as a substrate ^13, 43^ (Figure 3A). Lastly,from cluster 2, MplI has been suggested with longer acyl-CoA as physiological substrates ^13, 43^ (Figure 3A). EsaI was introduced as control, being ACP-dependent ^47^.

**Figure 3:**
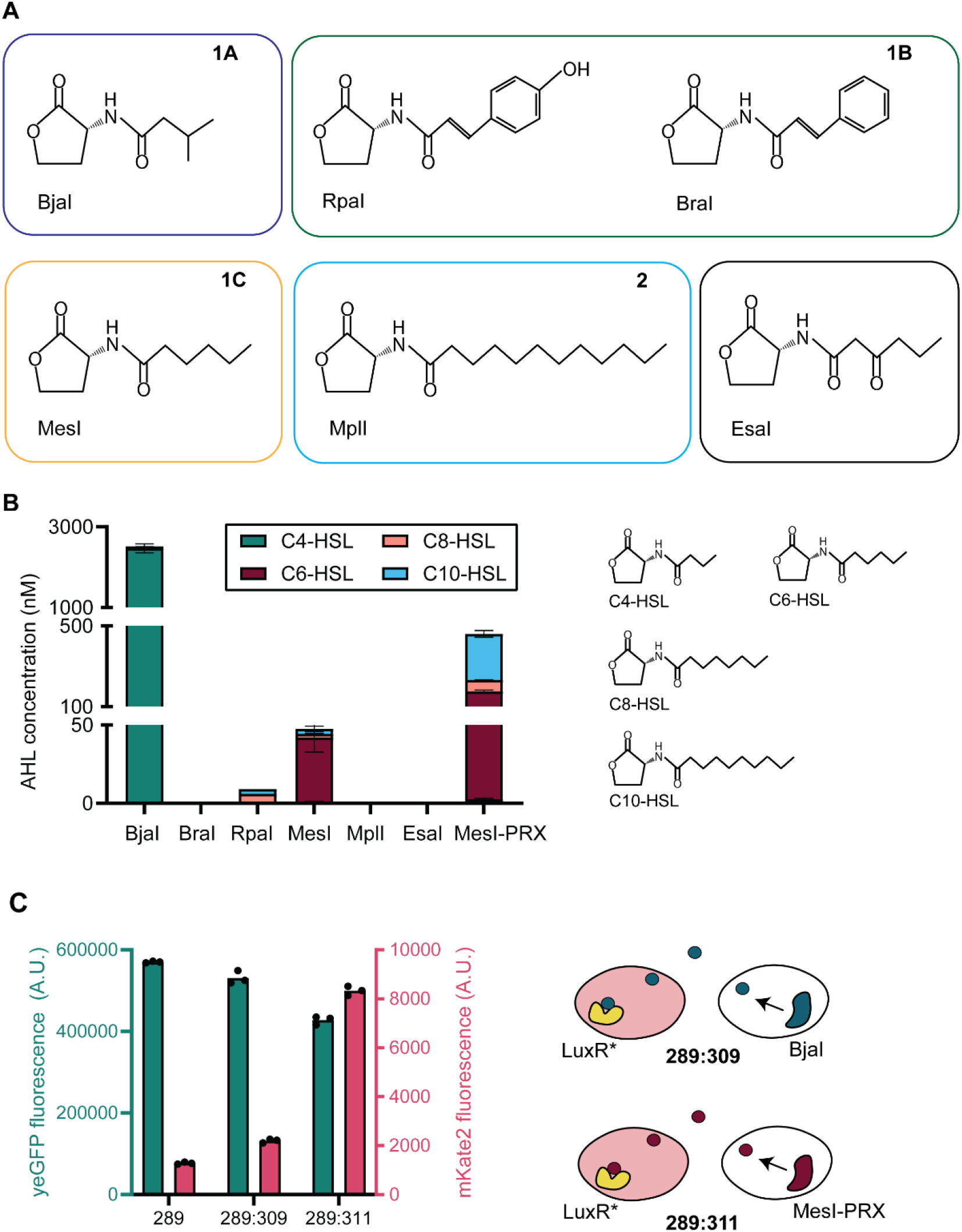
Different acyl-homoserine lactone synthases and AHL production. **(A)** overview of the different CoA-dependent clusters 1A, 1B, 1C and 2 ^40^ and the corresponding synthases chosen to introduce in yeast. EsaI is included as a control, which is an ACP-dependent synthase. Associated product of each synthase is shown. **(B)** Each of the indicated synthases is introduced into yeast strain ACA45 (5x*luxo*-*amdS*, 5x*luxo*-yeGFP, p*PGK1*-*GAL4*_AD_-*luxR*\*, *SAM2*-*MET6* (++)). Cells are grown in duplicate on fedbatch medium (EnPump) and supernatant is harvest after 24 h. AHL production is determined by LC-MS analysis and mean and standard deviation of the mean are shown. (**C**) yeGFP and mKate2 fluorescence of sensing strain AAA289 (5x*luxO*-mKate2, p*TEF1*_*luxO*-yeGFP, p*PGK1*-*GAL4*_AD_-*luxR\**) in monoculture and co-cultures of sensing strain AAA289 together with sending strains AAA309 (*SAM2*-*MET6, bjaI*) and AAA311 (*SAM2*-*MET6, mesI*-PRX) after 24 h on fedbatch medium. Individual points of biological replicates (n=3) are shown. A.U.: arbitrary units.

These synthases were expressed in combination with an additional copy of *MET6* and *SAM2*, which were previously shown to boost SAM levels ^48^ and modestly improve C8-HSL production in yeast ^27^. To assess AHL synthesis, we introduced the LuxR*-based biosensor as well (resulting in platform strain ACA045; *SAM2*-*MET6* (++), 5x*luxO*-yeGFP, 5xluxO-*amdS*, p*PGK1*-*GAL4*_AD_-*luxR*\*), since it exhibits the broadest AHL responsiveness (Figure 1E). Flow cytometric analysis of biosensor activation after 24 h on fed-batch medium revealed that BjaI, BraI, RpaI and MesI are able to trigger a fluorescent response (Figure S4). These 4 synthases can therefore be used in a QS-circuit in combination with LuxR*.

LC-MS analysis of the supernatant further assessed the distinct production profiles (Figure 3B). In accordance with its native substrate, MesI predominantly produces C6-HSL (41 ± 9 nM), with over tenfold lower concentrations of C8-HSL (2.4 ± 0.4 nM) and C10-HSL (3.2 ± 1.7 nM) produced as well. On the other hand, we found that BjaI predominantly produces C4-HSL with concentrations of 2.5 ± 0.1 µM, and C6-HSL as additional product at much lower concentrations of 37 ± 2 nM. Additionally, traces of C8-HSL (4 ± 0 nM) and C10-HSL (11 ± 0 nM) were detected as well. While the preferred substrate of BjaI is isovaleryl-CoA (IV-CoA), previous studies found that BjaI can act on butyryl-CoA as well, albeit with a 4.9 lower k_cat_/K_M_, compared to IV-CoA ^13^. Lastly, RpaI produces low amounts of C10-HSL (3 ± 0 nM) and C8-HSL (6 ± 0 nM). This indicates some flexibility in substrate utilization by these synthases, but might also reflect the differences in the physico-chemical conditions of the yeast cytosol. In line with the fluorescent characterization, no AHLs were detected for MplI and EsaI. For BraI however, we were not able to detect any AHLs using LC-MS either.

Since AHLs with longer chains become more and more hydrophobic, we postulated that some molecules could be missed when analysing directly from the supernatant. To test whether hydrophobic AHLs were retained intracellularly, we used an organic phase extraction. This time, we could detect C12-HSL (Figure S5), produced by BjaI, MesI and RpaI. However, these concentrations were below 1 nM, which precludes accurate estimation of the concentration. For C14-HSL (Figure S6), only BjaI produced a molecule matching this retention time, again at too low concentrations. Still, no AHLs were detected for BraI, which could indicate that the biosensor is able to pick up signals that LC-MS cannot or that the extracellular concentration of the produced AHLs is lower than the intracellular.

### Targeting synthases to peroxisomes

While acyl-CoA is present in the cytosol, concentration of acyl-CoA with different chain lengths are presumably higher inside the peroxisomes ^49^. Possibly, AHL production can be further increased by improving the availability of preferred CoA-substrate.Peroxisomes play a prominent role in β-oxidation and β-oxidation intermediates include different chain lengths of acyl-CoAs. Moreover, targeting enzymes to peroxisomes to enhance metabolites derived from β-oxidation intermediates have proven successful ^49^. Therefore, we evaluated AHL production when the synthases were targeted to the peroxisomes using a peroxisomal tag (PRX; LGRGRRSKL) ^50^. To be able to compare AHL-production profiles to the untagged synthases, strain ACA045 was used again as platform strain. Localization to the peroxisome increased the fluorescent response for BraI and MesI (Figure S4). Based on LC-MS analysis, MesI produces approximately 4-, 23 and 72-fold more C6-HSL, C8-HSL and C10-HSL respectively, compared to untagged MesI (Figure 3B). Similarly, production of C10-HSL was increased 10-fold for BjaI, when targeted to the peroxisome (Table S1). In contrast, C4-HSL production was reduced greatly (from 2468 ± 112 nM to 13 ± 0 nM) in the strain synthesizing BjaI-PRX, which could reflect a difference in availability of butyryl-CoA availability. The differences in AHL production between the tagged and untagged synthases might reflect a difference in substrate availability in the peroxisomes. Moreover, in previous studies, no activity was detected with octanoyl-CoA or decanoyl-CoA as substrate ^13^. But in our study we show moderate activity of BjaI on these substrates *in vivo* in yeast, which is enhanced by targeting the enzyme to the peroxisome. The results in this study suggest that BjaI can accept more substrates when expressed in yeast than previously thought from *in vitro* studies, which could be a reflection of differences in environment such as pH ^51^ or ion availability ^52^. However, we cannot exclude the possibility that the peroxisomal tag influenced the folding and therefore the activity of the enzyme.

For RpaI and BraI, there was no difference in the detected AHL production profile (Table S1). Specifically, for BraI-PRX, again we were not able to detect any specific signaling molecule produced. Using untargeted LC-MS we were not able to distinguish any signal for others HSLs either and the product of BraI in yeast therefore remains to be elucidated.

### Cross-feeding of AHL-molecules in co-cultures

Having now established multiple new QS-circuits in monoculture, we tested whether this communication can still occur when the modules are separated in sender and sensor strains. BjaI and MesI-PRX were tested as sender strains since these were the highest producers, and the dual purpose LuxR*-based system was used as sensor strain since this system can sense C4-HSL, C6-HSL and C10-HSL, the main products of BjaI and MesI-PRX. Strains were cultured together on fed-batch medium for 24 h and yeGFP and mKate2 fluorescence was analysed by flow cytometry and compared to AAA289 monoculture. The strain carrying BjaI elicited a limited response in the biosensor strain, with 7% repression of yeGFP and a 1.7-fold induction of mKate2 (Figure 3C). MesI-PRX on the other hand, could induce mKate2 up to 6.5-fold while repressing 25% of yeGFP (Figure 3C).

### Increasing substrate availability of hydroxycinnamoyl-CoAs

To engineer an orthogonal QS-couple, we need to pair another synthase with a biosensor that does not respond to the previously established MesI-PRX sender. RpaI natively utilizes *p*-coumaroyl-CoA to produce *p*-coumaroyl-HSL. Since coumaric acid and *p*-coumaroyl-CoA are not natively synthesized by *S. cerevisiae*, we introduced tyrosine ammonia-lyase from *Flavobacterium johnsoniaeu* (FjTAL) ^53^ and 4-coumarate:CoA ligase from *Arabidopsis thaliana* (At4CL) ^54^ to provide the necessary precursors (Figure 4A). To test substrate promiscuity, BjaI, BraI, MesI and EsaI were expressed with TAL and 4CL1 as well. Only co-expression of TAL, 4CL1 and RpaI led to high concentrations of *p*-coumaroyl-HSL of approximately 1.5 µM (Figure 4B).

**Figure 4:**
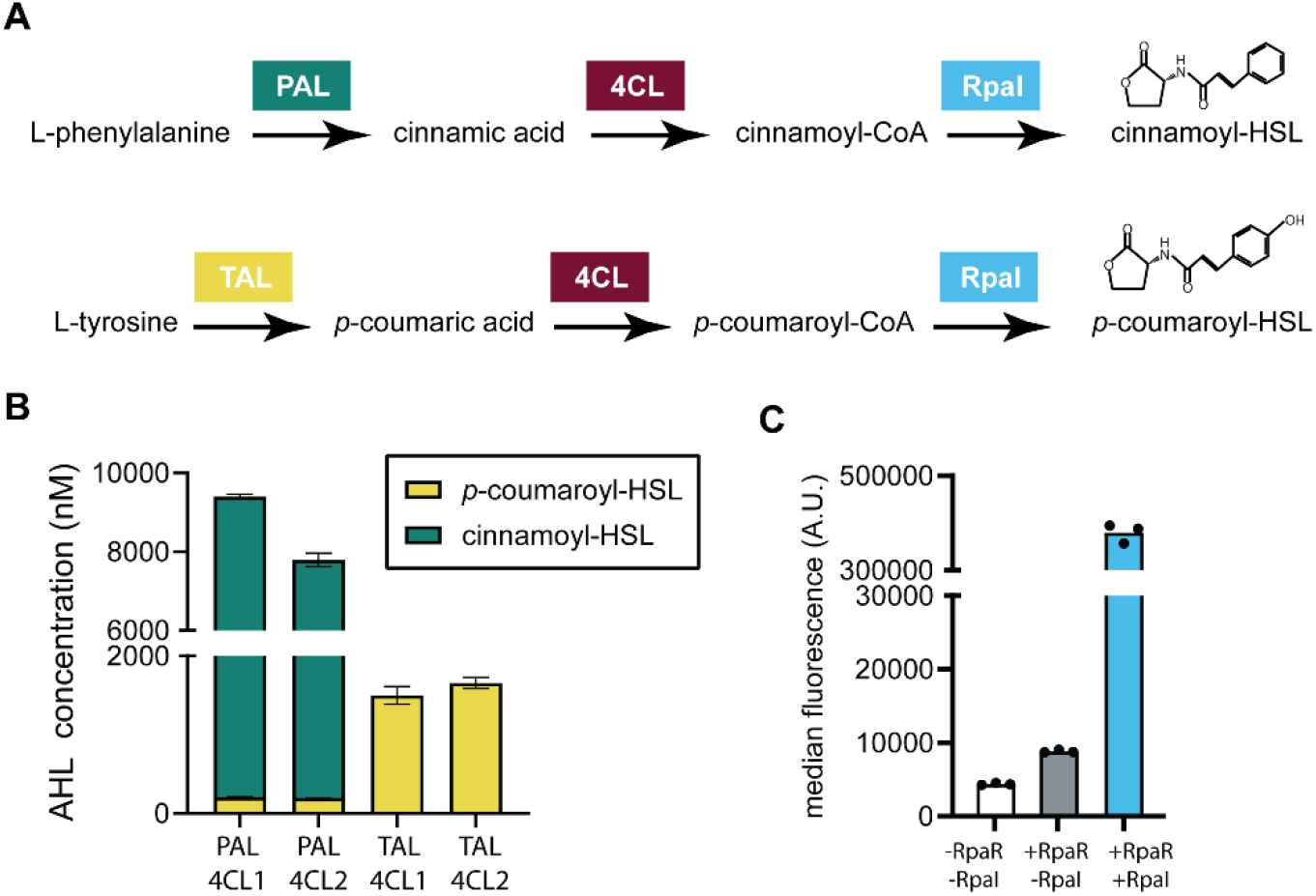
Engineering hydroxycinnamoyl-HSL production and QS circuit in yeast. **(A)** Overview of biosynthetic pathway and the introduced enzymes for the production of cinnamoyl-HSL and *p*-coumaroyl-HSL in yeast. **(B)** Analysis of cinnamoyl-HSL and *p*-coumaroyl-HSL by strains AAA119 (p*PGI1*-AtPAL, p*ACT1*-At4CL1, p*TDH3*-*rpaI*), AAA121 (p*PGI1*-AtPAL, p*ACT1*-At4CL2, p*TDH3*-*rpaI*), AAA092 (p*PGI1*-FjTAL, p*ACT1*-At4CL1, p*TDH3*-*rpaI*) and AAA123 (p*PGI1*-FjTAL, p*ACT1*-At4CL2, p*TDH3*-*rpaI*). Cells are grown in duplicate on fedbatch medium (EnPump) and supernatant is harvested after 24 h. AHL production is determined by LC-MS analysis and mean and standard deviation of the mean are shown. The concentration of *p*-coumaroyl-HSL was quantified using an external calibration standard, while cinnamoyl-HSL concentration was estimated assuming an equivalent MS response to *p*-coumaroyl-HSL. **(C)** yeGFP fluorescence of strains strains AAA160 (-RpaR -RpaI: 5x*rpaO*-yeGFP), AAA258 (+RpaR -RpaI: 5x*rpaO*-yeGFP, p*PGK1*-VP48-*rpaR*) and AAA263 (+RpaR +RpaI: 5x*rpaO*-yeGFP, p*PGK1*-VP48-*rpaR*, p*PGI1*-FjTAL, p*ACT1*-4CL2, p*TDH3*-*rpaI)* after 24 h on fedbatch medium. Individual points of biological replicates (n=3) are shown. A.U.: arbitrary units.

We further extended the AHL production repertoire in *S. cerevisiae* by introducing a metabolic pathway for the synthesis of cinnamoyl-CoA production as well, via phenyl-alanine ammonia-lyase (AtPAL) and 4CL (Figure 4A). LC-MS analysis revealed substantial accumulation of cinnamoyl-HSL species, but since we did not have a standard, we could only estimate the concentration of cinnamoyl-HSL (Figure 4B, Figure S7). Additionally, we tested two variants for 4CL (At4CL1 and At4CL2), together with PAL and TAL as well. We found that 4CL2 was slightly better in combination with TAL (AAA123) while 4CL1 was better in combination with PAL (AAA119), in line with a previous study in which 4CL2 was used for the production of *p*-coumaroyl-CoA ^54^ and 4CL1 was used in combination with PAL ^55^. We analysed the growth of the highest *p*-coumaroyl-HSL producing strain (AAA123) on SC medium in shakeflasks and measured a modest 9% slower growth rate of 0.35 ± 0.00 h^-1^ compared to a non-engineered yeast strain which displayed a growth rate of 0.38 ± 0.01 h^-1^.

### Engineering a p-coumaroyl-HSL-based QS control loop

We introduced our RpaR-based *p*-coumaroyl-HSL biosensor into our *p*-coumaroyl-HSL producing strain AAA123 and analysed fluorescence after growing on fedbatch medium. The yeGFP fluorescence level was induced 43-fold compared to a non-producing strain biosensor strain (Figure 4C). This shows that the *p*-coumaroyl-HSL produced can be sensed by the RpaR-based biosensor, constituting a novel RpaI-RpaR QS circuit.

### Establishing orthogonal QS couples

For the development of orthogonal QS pairs, it is essential to decouple signal production and signal sensing into separate yeast strains, such that each strain mainly produces one QS molecule while sensing only the main signal produced by the sender strain. Building on our previously established co-culture communication using the MesI–PRX production module and LuxR* (Figure 3C), and our orthogonal RpaI-RpaR pair, we constructed producer–sensor strains by combining RpaR with MesI–PRX and LuxR with RpaI. To minimize unintended cross-activation by less abundant AHLs produced by RpaI (Figure 3B), we started with LuxR_N86K_ instead of LuxR*, since N86K alone confers slightly less sensitivity ^27^. In these strains, LuxR_N86K_ controlled expression of mKate2 (AAA312) and RpaR of yeGFP (AAA320) and fluorescence was measured after 24 h on fedbatch medium. While RpaR-based reporter expression was observed when RpaI was co-expressed within the same strain (Figure 4C), no activation was detected when RpaI was expressed in a separate strain in co-culture (Figure 5A). Moreover, LuxR_N86K_ was not sensitive enough to sense the production of AHL by MesI-PRX (Figure 5A).

**Figure 5:**
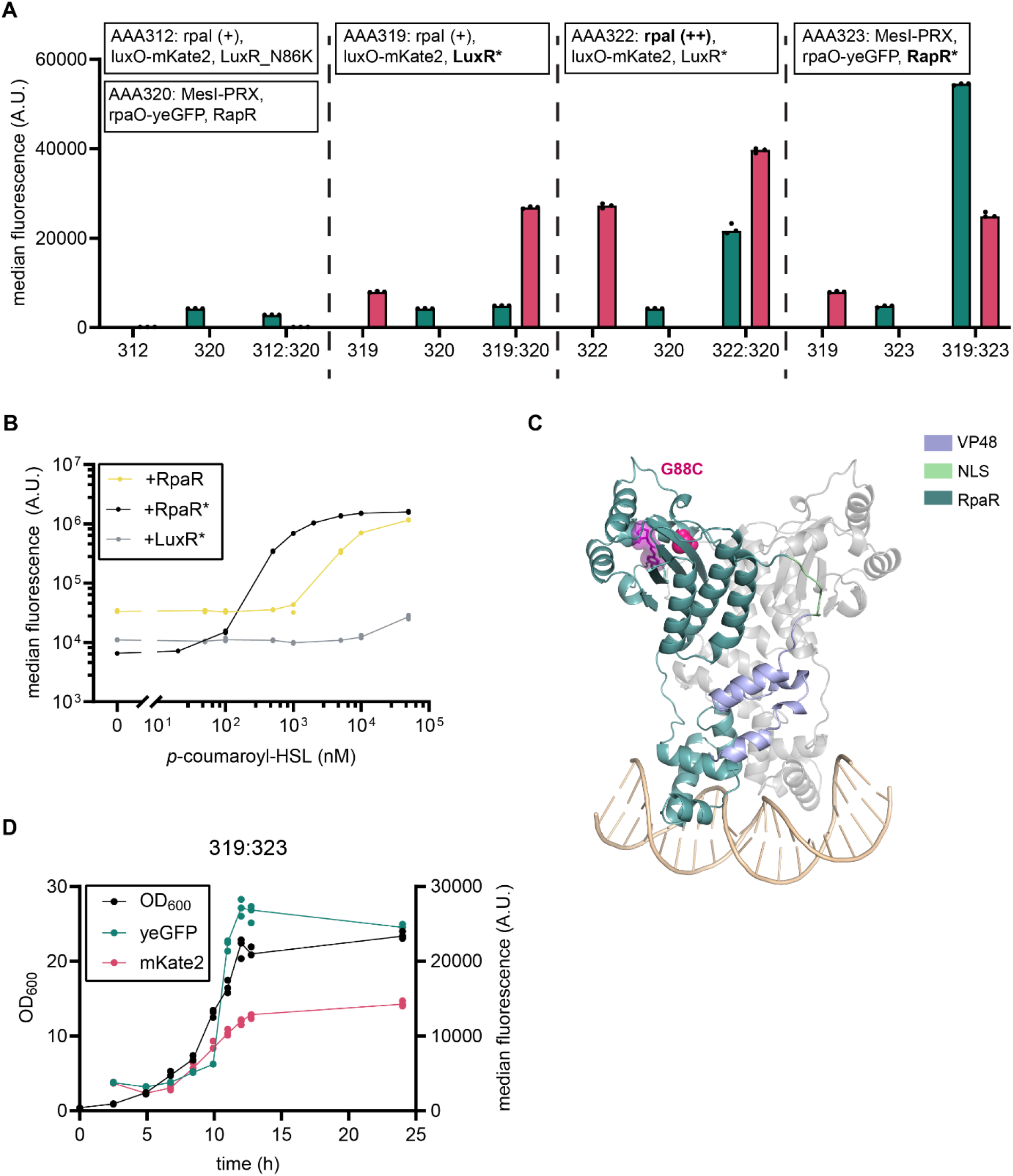
Construction of orthogonal QS-pairs. **(A)** yeGFP and mKate2 fluorescence of strains AAA312 (TAL-4CL2-*rpaI*, 5x*luxO*-mKate2-*luxR*_N86K_) and AAA320 (SAM2-MET6 (++)-*mesI*-PRX, 5x*rpaO*-yeGFP-*rpaR*), AAA319 (TAL-4CL2-*rpaI*, 5x*luxO*-mKate2-*luxR*\*) and AAA320, AAA322 (TAL-4CL2 (++)-*rpaI*, 5x*luxO*-mKate2-*luxR*\*) and AAA320, AAA319 and AAA323 (SAM2-MET6 (++)-*mesI*-PRX, 5x*rpaO*-yeGFP-*rpaR*\*), in monoculture and co-cultures after 24 h on fedbatch medium. **(B)** Dose-response curves of yeGFP fluorescence levels 6 h following supplementation of 0–100,000 nM *p*-coumaroyl-HSL, tested for strains AAA156 (+LuxR*: 5x*luxO*-yeGFP, p*PGK1*-*GAL4*_AD_-*luxR\**), AAA258 (+RpaR: 5x*rpaO*-yeGFP, VP48-*rpaR*) and AAA312 (+RpaR*: 5x*rpaO*-yeGFP, p*PGK1*-VP48-*rpaR*\*). **(C)** Structural mapping of mutations identified in RpaR*. The structural model was generated using Boltz-2 ^39^ and includes a RpaR* dimer co-folded with the ligand *p*-coumaroyl-HSL (purple) and the DNA operator *rpaO* (yellow). **(D)** Growth and median fluorescence monitored during aerobic growth on fedbatch medium in shakeflasks of co-culture AAA319 and AAA323. For A, B, D: Individual points of biological replicates (n=3) are shown. A.U.: arbitrary units.

We next replaced LuxR_N86K_ with the more sensitive LuxR* variant (AAA319) and mKate2 fluorescence was increased 3.4-fold when co-culturing with MesI-PRX-RpaR (AAA320). To further improve *p*-coumaroyl-HSL production, we changed the promoters of TAL and 4CL2 to p*PGK1* and p*TEF1* (AAA322). This increased the self-induction of mKate2 by LuxR* 3.4-fold. Consistent with this observation, extending dose–response measurements to higher ligand concentrations revealed that LuxR* exhibits modest activation at extracellular *p*-coumaroyl-HSL concentrations approaching 100 µM (Figure 5B). Additional induction of mKate2 caused by co-culturing with MesI-PRX was only 1.5-fold. On the other hand, RpaR now showed a 5-fold increase in yeGFP fluorescence when co-cultured with the high-producing RpaI module (AAA322).

### Increasing sensitivity of RpaR

As increasing *p*-coumaroyl-HSL production improved intercellular signaling at the cost of increased self-activation, we next sought to enhance system performance by increasing the sensitivity of RpaR. We therefore constructed a mutant library and performed the counter selection with 400 nM of C10-HSL, to prevent self-activation by MesI-PRX, and the selection with 1 µM of *p*-coumaroyl-HSL. The selected population displayed a small subpopulation with a more fluorescent phenotype (Figure S8) and 48 colonies were tested to find single cells with an improved phenotype. One colony was selected (ACA083) and sequenced and the variant was used to create an improved RpaR-based biosensor (AAA321; 5x*rpaO*-yeGFP, VP48-*rpaR*_G136C_), which will be referred to as RpaR* in this study. The performance of RpaR* mutant was evaluated by dose– response analysis and revealed a lower basal expression level as well as higher sensitivity and broader operational range (Figure 5B).

Two mutations were identified in RpaR, of which only one non-synonymous; G88C. This mutation resides in the ligand binding pocket although it does not appear to interact directly with the ligand in the generated Boltz2 structural model ^39^ (Figure 5C, Figure S9). Such a bond could potentially constrain local pocket flexibility and stabilize a ligand-responsive state.

### Establishing intercellular communication in yeast co-cultures

Finally, we combined this more sensitive RpaR* with MesI-PRX (AAA323) and paired it with the original lower *p*-coumaroyl-HSL producing module and LuxR* (AAA319). Co-culturing both strains led to 5.2-fold increase in yeGFP and 3.1-fold increase in mKate2, showing bidirectional QS-dependent communication.

To get an idea of the timing of induction and growth performance of the strains, we also tracked its growth and fluorescence per cell using flow cytometry on fed batch medium. This indicated delayed induction for both strains as well as fast growth with a growth rate of 0.35 ± 0.00 h^-1^. LuxR* mediated induction was estimated after approximately 8 hours, and RpaR* mediated induction after almost 10 hours, with final mKate2 induction level of 3.9-fold and yeGFP induction of 6.4-fold.

As AHL and *p*-coumaroyl-HSL increases with biomass concentration, final induction of its partner can be further increased through longer fedbatch cultivation. Alternatively, the dual-purpose design (Figure 2F) could potentially be used to keep induction levels stable throughout the cultivation, by coupling synthase expression to the repressive module.

## Conclusion

In this work, we expanded the current quorum-sensing toolkit in *Saccharomyces cerevisiae*, by engineering of QS synthases and cognate biosensors, and demonstrated their functionality and orthogonality in co-culture. Our results confirm that QS-mediated intercellular communication can be robustly reconstituted in a eukaryotic host. We identified the G136C mutation in RpaR as a key determinant of increased sensitivity,while F32Y was suggested to be at least partially linked to the improved dynamic range and sensitivity of TraR. Furthermore, peroxisomal targeting significantly increased C10-HSL production in combination with MesI, and expression of 4CL2, TAL and *rpaI* enabled production of *p-*coumaroyl-HSL. We identified MesI-PRX-LuxR and RpaI-RpaR as functional orthogonal QS-pairs. The observed 3.9-fold and 6.4-fold induction in co-cultures relative to monocultures confirms that reporter activation is driven by QS-mediated intercellular signaling rather than cell-autonomous expression or environmental effects.

Since AHL-based communication systems are already well established in bacterial cultures ^15-21^, expanding quorum-sensing tools in yeast enables their direct integration into coordinated interspecies systems ^56^. Such communication strategies are particularly attractive for metabolic engineering applications, where one strain produces a toxic or metabolically burdensome intermediate that is subsequently converted into a final product by a second strain. QS-based induction allows production of these intermediates to be delayed until sufficient cell density is reached, thereby reducing metabolic burden during early growth ^57^. Moreover, separating sender and sensor modules across different strains prevents premature activation in pre-cultured monocultures, ensuring that toxic intermediates are not produced in the absence of their co-culture counterpart ^58^. Beyond metabolic engineering, AHL-based cell–cell communication provides a framework for coordinating collective behaviors at defined population densities. For example, this system could be used to implement density-dependent induction of flocculation ^59^ or biofilm formation ^60^, again with the benefit of separation of sender and sensor modules to prevent premature activation during pre-cultivation.

## Materials and methods

### Strains and growth media

The *Saccharomyces cerevisiae* strains used in this study are listed in Table S2 and are derived from CEN.PK110-10C (MAT-a *URA3 LEU2 TRP1 his3*) and CEN.PK2-1C (MAT-a*ura3 his3 leu2 trp1*). For experimental assays, the engineered yeast strains were cultivated in synthetic complete medium (6.7 g L^-1^ yeast nitrogen base without amino acids, 1.62 g L^-1^ yeast synthetic drop-out medium supplement without leucine, 0.2 g L^-1^ leucine, 20 g L^-1^ glucose, pH set to 5.6 with 2M KOH). During strain construction, the engineered yeast strains were routinely cultivated at 30°C in synthetic complete medium lacking histidine (6.7 g L^-1^ yeast nitrogen base without amino acids, 1.92 g L^-1^ yeast synthetic drop-out medium supplement without histidine supplemented with 20 g L^-1^ glucose, pH set to 5.6 with 2M KOH). For selection using clonNAT marker, synthetic media lacking histidine or lacking histidine and uracil and containing monosodium glutamate (SMG) was used (1.7 g L^-1^ yeast nitrogen base without amino acids and ammonium sulfate, 1 g L^-1^ monosodium glutamate, 1.92 g L^-1^ yeast synthetic drop-out medium supplement without histidine or 1.39 g L^−^1 yeast synthetic drop-out medium supplement without histidine, leucine, tryptophan and uracil supplemented with 0.2 g L^−^1 leucine and 0.07 g L^−^1 tryptophan, 20 g L^-1^ glucose, pH set to 5.6 with 2M KOH, 2% (w/v) agar in case of plates) supplemented with 100 mg L^-1^ nourseothricin. Extra-buffered fed-batch medium contained 5.0 g L^−1^ KH_2_PO_4_, 0.5 g L^−1^ MgSO_4_·7H_2_O, 14.4 g L^−1^ (NH_4_)_2_SO_4_, 1 ml L^-1^ trace elements, 1 ml L^-1^ vitamins, 0.125 g L^-1^ histidine, 0.5 g L^-1^ leucine, 0.075 g L^-1^ tryptophan, 0.15 g L^-1^ uracil, 20 g L^-1^ glucose and 40 g L^-1^ EnPump 200 substrate (Enpresso, Berlin, Germany). Fed-batch was started with 8 mL L^-1^ of enzyme mix. Synthetic minimal medium (SMD) contained 3.0 g L^−1^ KH_2_PO_4_, 0.5 g L^−1^ MgSO_4_·7H_2_O,5.0 g L^−1^ (NH_4_)_2_SO_4_, 1 ml L^−1^ trace elements and vitamins ^61^ supplemented with 20 g L^-1^ glucose. Counterselection medium was prepared by adding fluoroacetamide to SMD to a final concentration of 20 g L^-1^. For selection medium, (NH_4_)_2_SO_4_ was replaced by 6.6 g L^−1^ K_2_SO_4_ and 0.6 g L^−1^ filter-sterilized acetamide.

For cloning and plasmid propagation, *Escherichia coli* strain DH5 was used, in Luria-Bertanii (LB) medium containing 100 µg mL^-1^ ampicillin.

### Plasmid construction

The plasmids used in the study are listed in Table S3. Codon-optimized versions of *rpaI, braI, bjaI, mesI* and *mplI* were synthesized by GeneArt Thermo Scientific (Table S6).ORFs for EsaR ^34^, EsaR_V220A_^34^, EsaR_D91G_^34^, RpaR ^20^, TraR ^20^ and LasR ^20^ were obtained from Addgene (#47660, #47645, #47646, #85148, #85150, #85147). Plasmid construction was performed by USER-cloning protocol, using fragment-specific primers (Table S5) and Phusion U high-fidelity DNA polymerase (New England Biolabs) according to manufacturer’s instructions. For construction of single ORF cassettes, the backbone was PCR-amplified with MAD3/MAD4 as described previously ^62^. For construction of plasmids with two ORFs, the backbone was linearized using SfaAI FD and BsmI FD (Thermo Scientific) ^63^. The PCR-amplified backbone, promoter, ORF, terminator, were added in equimolar amounts with cutsmart buffer and USER-enzyme and handled according to supplier’s protocol. 2 µL of this mixture was used in the subsequent transformation with *E. coli* and the whole mixture was plated on selective medium.

### Yeast strain and library construction

Yeast strains ACA001 and AAA001 were used for all yeast transformations, which contained a plasmid carrying Cas9, using HIS3 as selection marker. Strain construction was performed using CRISPR-Cas9-mediated genome editing, enabled by introduction of one or two gRNA-plasmid and one or two NotI-digested plasmids encoding the integration cassette following EasyClone protocol ^63^ and as described in Table S2.Exceptions include the integration of p*PGK1*-*luxR*_N86K_-tADH1 into X-4, which was introduced as two linear PCR-amplified fragments (Table S4) introducing the N86K mutation and assembled using homologous recombination into the yeast genome. Moreover, for yeast library construction, DNA-fragments were prepared as described previously ^27^: Error-prone PCR (epPCR) was performed on 750 ng template DNA of plasmids pAvA169 and pAvA171, for RpaR and TraR respectively, using the Agilent GeneMorph mutagenesis II kit. Primers AA73/AA7 were used for one round of error-prone PCR. Subsequent PCR was performed with this PCR product as template, using primers AA74/AA78. Homology arms for integration in X-4 were amplified using AA757/AA76 and AA79/AA80 and these fragments contained the promoter sequence or the terminator sequence, respectively. Yeast cells were transformed using a heat-shock transformation protocol ^64^, with the following adaptations: Cells were washed in 0.1 M LiAc prior to addition of the transformation mixture. After resuspending the cells in the transformation mixture, the cells were incubated for 15 minutes at 30°C, prior to 30 minutes heat-shock at 42°C. Transformations were plated on SMG -HIS + NAT or SMG - HIS -URA + NAT. For library construction, 400 mL yeast culture of OD_600_ = 2 was divided over 10-16 tubes and the biomass was used in separate transformations. The total biomass was pooled together before the recovery step and after recovery, all cells were added to 50 mL SMG -HIS + clonNAT. By plating for single colonies on SMG -HIS + clonNAT right after the transformation, it was determined that the RpaR-library contained 7.6 * 10^5^ possible variants and the TraR-library 1.3 * 10^6^. For transformations, colonies were restreaked once on selective medium and twice on non-selective medium and correct integration was confirmed by colony PCR using RedTaq MM. Removal of gRNA plasmid carrying a clonNAT marker was confirmed via restreaking on SC -HIS and SMG -HIS + NAT. For removal of plasmids with auxotrophic markers,colonies were grown overnight on non-selective liquid medium before dilutions were plated on YPD to obtain single colonies. Verification of plasmid removal was performed by transferring all colonies to filter paper (Whatman, grade 1 round filter paper, 150 mm) and subsequently stamping these colonies on new plates on YPD and on each individual selection (i.e., SC-HIS, SC-URA). Correct colonies were inoculated in SC or SC -HIS and grown till late exponential phase after which glycerol was added up to 30% and cells were stored at -70°C.

### Flow cytometric analysis

A 1 mL aliquot of glycerol -70°C freezer stock was used to inoculate 50 mL of SC medium (pH 5.6) and grown for 16 hours. For biosensor characterization, the culture was diluted 1:20 in 200 µL SC medum ± inducer in 96-deep-well culture plates and grown for 6 hours. Cells were washed and diluted 1:4 in PBS prior to analysis by flow cytometer. 20,000 events were recorded for each well, For the other experimental assays, the culture was diluted 1:10 in 1 mL SC medium and grown for 8 hours. After 8 hours, 100 µL of this culture was transferred to 4 mL fed batch medium in 50 mL CELLSTAR® CELLREACTOR tubes (Greiner Bio-One). After 24 h cells were washed and diluted 1:10 in PBS prior to analysis by flow cytometer, while the supernatant was stored at -20°C for subsequent analysis and/or extraction. Flow cytometry analysis was performed on the Novocyte. 30,000-50,000 events were recorded for each well. The threshold for event detection was at >150,000 FSC-H with a core diameter of 10.1 µm. For yeGFP, excitation was performed with a blue laser (488 nm) and emission detection with a 530/30 nm BP filter. For mKate2, excitation was performed at 561 nm and emission detection with a 615/20 nm BP filter. FSC-A was plotted against FSC-H to gate for singlets events (Figure S10). Gated events were used to determine the median fluorescence of the population. For co-culture samples, events were gated based on forward scatter height (FSC-H) and fluorescence intensity to identify yeGFP- and mKate2-expressing populations. Green (yeGFP) and red (mKate2) fluorescence channels were plotted against FSC-H, and gates were drawn around each fluorescent population (Figure S11). The median fluorescence intensity of yeGFP and mKate2 was then calculated for each gated population using FlowJo. the fluorescence Analysis of flow cytometric data was performed using FlowJo licensed software.

### AHL extraction

Supernatant (100 µL) was extracted with an equivalent volume of ethyl acetate. The mixture was shaken on a vortex for 30 seconds and the phases were allowed to separate. The top layer was collected in a new tube, and the extraction process was repeated one more time. Both extracts were pooled together and were evaporated using Vacufuge plus (Eppendorf). The dried extracts were stored at -20°C till detection. Before analysis, they were dissolved in HPLC-grade methanol and filtered.

### LC-MS/MS analysis

#### By TripleQuad LC-MS/MS

Detection and quantification of acylated homoserine lactones from yeast supernatant samples and extracted samples was performed as described previously ^27^. Briefly,chromatography was performed on a 1290 Infinity II UHPLC system (Agilent Technologies). Separation was achieved on a Kinetex XB-C18 column (100 × 2.1 mm, 1.7 µm, 100 Å, Phenomenex, Torrance, CA, USA), with formic acid (0.05%, v/v) in water and acetonitrile (supplied with 0.05% formic acid, v/v) as mobile phases A and B respectively. The flow rate of the mobile phase was 400 µL min^-1^ and the column was maintained at 40 °C. The liquid chromatography was coupled to an Ultivo Triplequadrupole mass spectrometer (Agilent Technologies) equipped with a Jetstream electrospray ion source (ESI) operated in positive ion mode. The ion spray voltage was set to 3000 V, with the dry gas flow at 10 L min^-1^ maintained at 325 °C. For the sheath gas a flow rate of 12 L min^-1^ was used at 400 °C. Nebulizing gas was set to 45 psi. Nitrogen was used as dry gas, nebulizing gas and collision gas. Precursor ion →fragment ion transitions were monitored using Multiple reaction monitoring (MRM), determined by direct infusion experiments of reference standards. Mass Hunter Quantitation Analysis for QQQ software (Version 10, Agilent Technologies) was used for data processing. Linearity in ionization efficiency was verified by analyzing dilution series that were also used for quantification of HSL in the samples.

#### By Quadrupole-time-of-flight (Q-TOF) LC-MS/MS

Q-TOF analysis was performed using a 1290 Infinity II UHPLC system (Agilent Technologies) equipped with Zorbax Eclipse XDB-C18 column (100 × 3.0 mm, 1.8µm, Agilent Technologies) as described previously ^27^. Formic acid (0.05%, v/v) in water was used as mobile phase A and acetonitrile containing 0.05% formic acid (v/v) was used as mobile phase B, at a flow rate of 400 µl min^-1^. The column temperature was maintained at 30 °C. The UHPLC system was coupled to a Bruker timsToF Pro mass spectrometer (Bruker, Bremen, Germany) equipped with an electrospray ioninazation source (ESI) operated in positive ion mode. The capillary voltage was set to +4200 V, the dry gas temperature to 200 °C, and the dry gas flow to 8 L min^−1^. Nitrogen was used as the dry gas, nebulizing gas, and collision gas. The nebulizing gas was set to 2.5 bar and collision energy to 10 eV. MS spectra were acquired over an *m/z* range of 50-1500 and MS/MS spectra over the same range, with a sampling rate of 5 Hz. Sodium formate clusters were used for mass calibration and all data files were calibrated by postprocessing.

### Mutational analysis

For comparative analysis across LuxR-family regulators, residues outside the activation domain were renumbered starting at 1, according to the full-length TraR and RpaR sequence. The structural models of VP48-NLS-TraR dimerized and bound to 3-oxoC8-HSL and *traO*, and the structure of VP48-NLS-RpaR bound to *p*-coumaroyl-HSL and *rpaO* were generated using Boltz-2 ^39^. Ligand binding mechanism was analyzed using PoseView ^65^. Figures were created using PyMol.

## Data availability

All data shown in figures are available in the Source data provided with this paper. There are no restrictions on data availability.

## Acknowledgements

This study is part of the project *Orthogonal quorum-sensing systems in yeast cell factories* with file number 019.231EN.007 of the Rubicon research programme which is financed by the Dutch research council (NWO) and awarded to AA. MH is funded by Novo Nordisk Foundation, grant number NNF22SA0078231 (Copenhagen Bioscience PhD Programme). This project has received funding from the Novo Nordisk Foundation, grant number NNF20CC0035580.

## Author contributions

**AA:** Conceptualization, Funding acquisition, Investigation, Visualization, Formal analysis, Methodology, Project administration, Writing – original draft, Writing – review & editing. **MH:** Formal analysis, Visualization, Writing – review & editing. **CC:** Resources, Investigation **MP:** Resources, Investigation. **EDJ:** Resources, Writing – review & editing. **MKJ:** Supervision, Writing – review & editing

All authors reviewed and approved the final version of the manuscript.

