## Supporting information for "Engineering orthogonal quorum sensing circuits using LuxR-type systems in yeast consortia"

#correspondence:

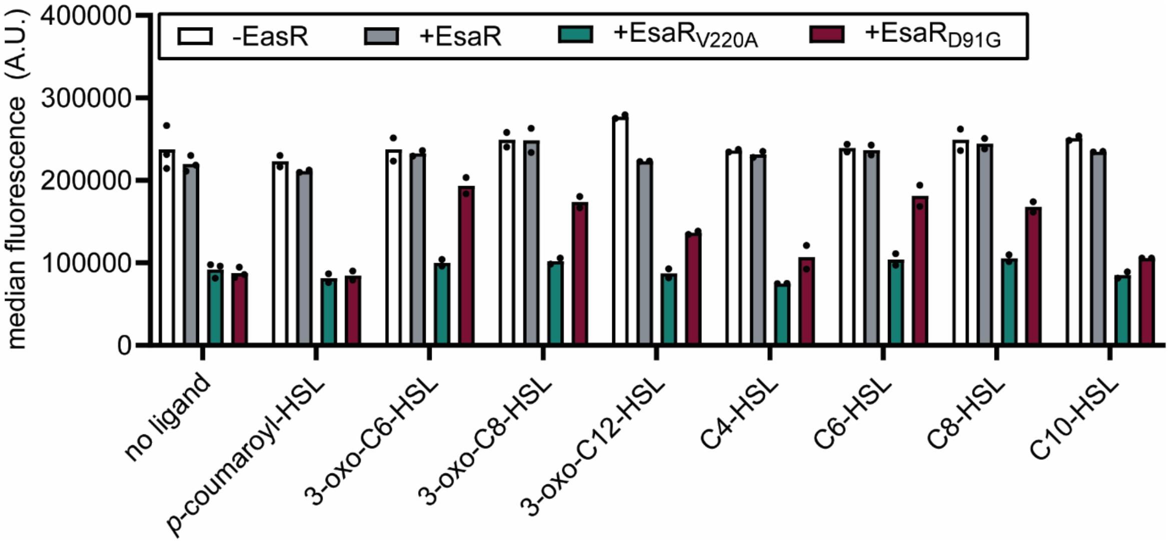

**Figure S1: yeGFP fluorescence levels in *EsaR*-based biosensor strains in response to addition of different AHL.** Median fluorescence was determined by flow cytometry for strains AAA015 (-*EsaR*; p*TEF1_esaO1*-yeGFP), AAA033 (+EsaR; p*TEF1_esaO*-yeGFP, p*PGK1*-*esaR*), AAA031 (+EsaR$_{V220A}$; p*TEF1_esaO*-yeGFP, p*PGK1*-*esaR*$_{V220A}$) and AAA032 (+EsaR$_{D91G}$; p*TEF1_esaO*-yeGFP, p*PGK1*-*esaR*$_{D91G}$) 6 h after supplementation with 100 μM of indicated ligand. Individual points of biological replicates (n=2) are shown. A.U.: arbitrary units.

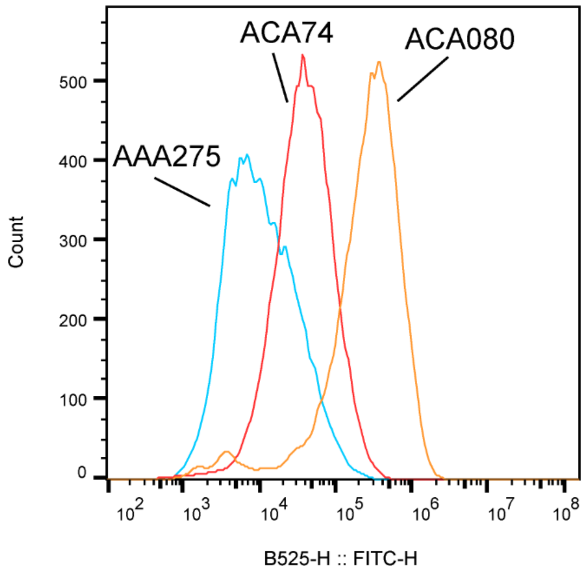

**Figure S2: Population fluorescence of VP48-*traR* library as determined by flow cytometry.** Using ep-PCR, a library was created of p*PGK1*-VP48-*traR* and this library was integrated into a yeast strain containing *amdS* and yeGFP under control of p*GAL*_core_5x*traO*. The yeast library was grown for 2 days on transformation selection media (SC -HIS +NAT), and transferred one more time for overnight growth on fresh transformation selection medium. Next, the population of selected transformants was transferred to counter-selection media (SMD + F-Ac) and grown for 1 day. Subsequently, cells were transferred twice on selection media (SMD -N + acetamide), containing 1 μM 3-oxo-C8-HSL. This final enriched population (ACA74) together with the parental biosensor strain (AAA275) and a selected colony from the library population, were grown in the presence of 1 μM 3-oxo-C8-HSL and analysed after 6 h on the flow cytometer.

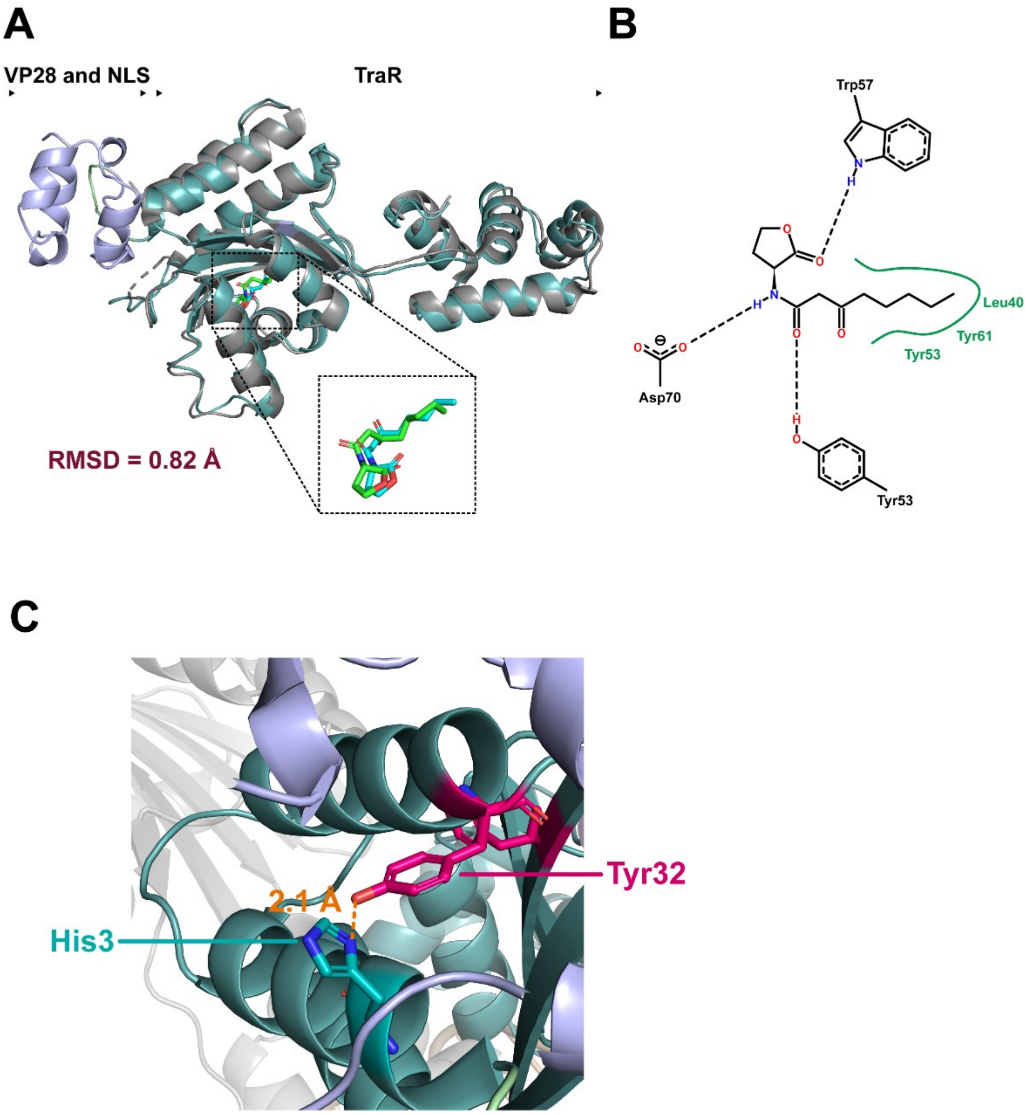

**Figure S3: Structural analysis of TraR*. (A)** VP48-NLS-TraR* Boltz-2 [1] structural model aligned with WT TraR crystal structure (PDB 1H0M). Ligand positioning in the ligand biding pocket and RMSD between TraR domains are shown. **(B)** Common ligand biding mechanism for 3-oxo-C8-HSL in the TraR crystal structure and TraR* structural model determined using PoseView**. (C)** Close up view of new hydrogen bond formed by the Phe32Tyr mutation with His3.

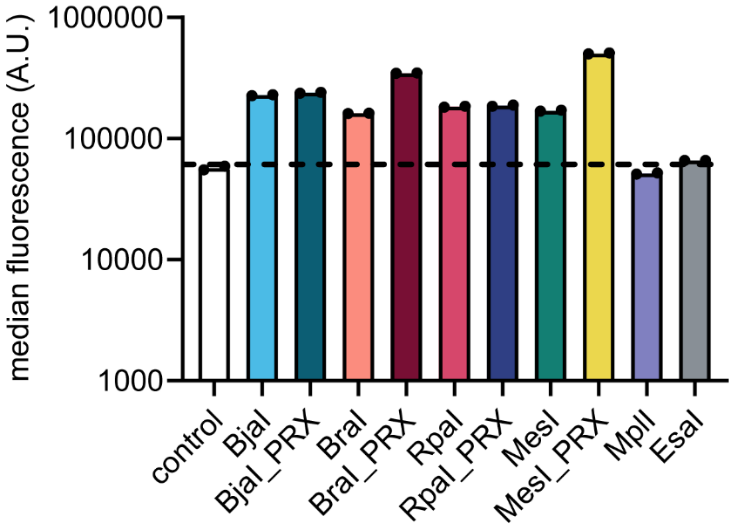

**Figure S4: yeGFP fluorescence induction in response to AHL production.** Each of the indicated synthases are introduced into strain ACA045 (control; *SAM2-MET6* (++), 5x*luxO*-yeGFP, 5x*luxO-amdS*, *luxR**) which contains p*PGK1*-*GAL4*$_{AD}$-*luxR*_gen1 [2] responsive to a wide range of AHLs. Increase of yeGFP fluorescence compared to the control strain therefore indicates self-sensing of the produced AHL by the indicated synthase. Strains ACA045 (control), ACA046 (BjaI), ACA071 (BjaI-PRX), ACA047 (BraI) ACA072 (BraI-PRX), ACA050 (RpaI), ACA070 (RpaI-PRX), ACA055 (MesI), ACA066 (MesI-PRX), ACA056 (MplI) and ACA049 (EsaI) were grown for 24 h on fedbatch medium and yeGFP fluorescence was analysed by flow cytometry. Individual points of biological replicates (n=2) are shown. A.U.: arbitrary units.

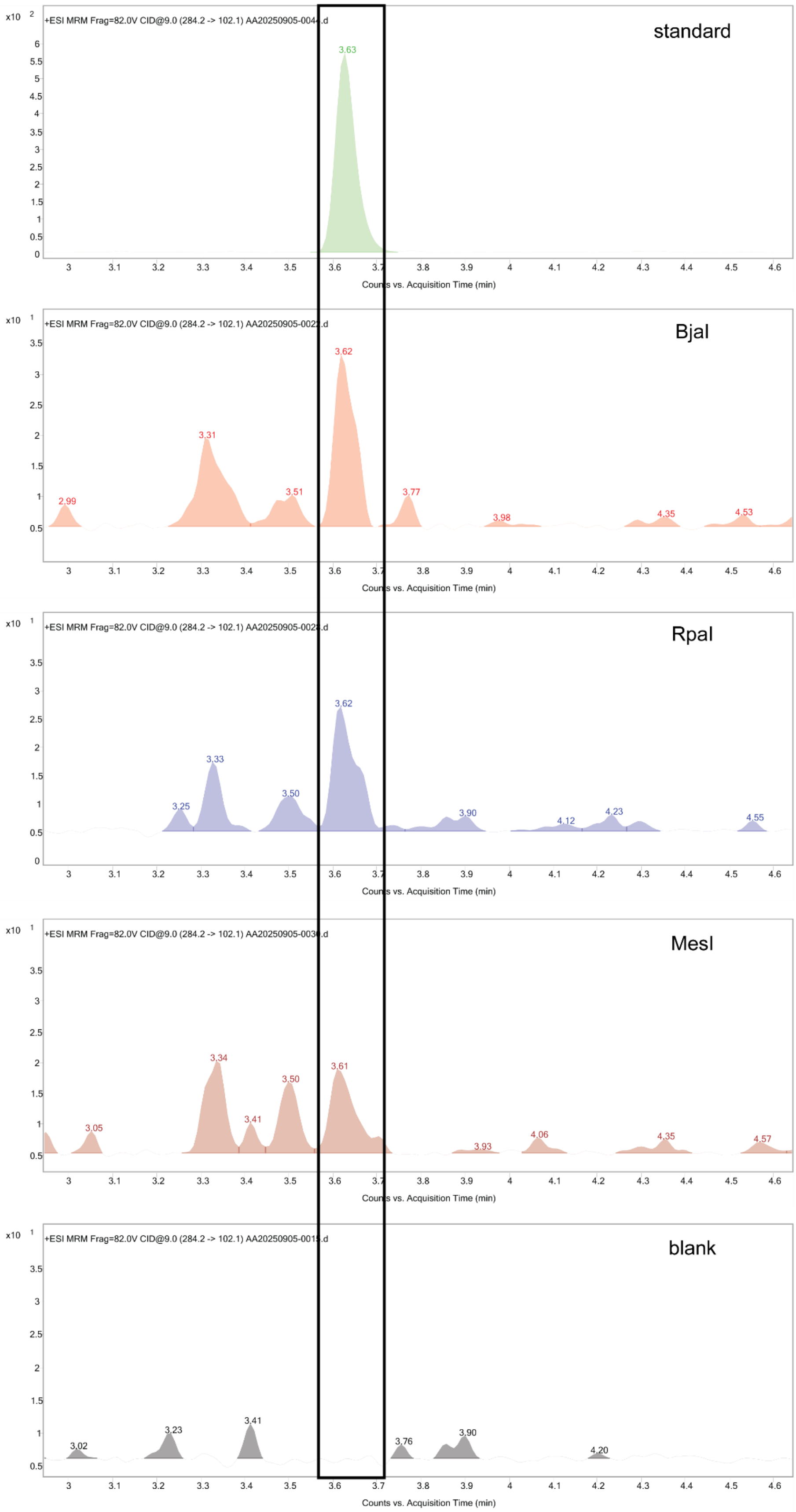

**Figure S5: Production of C12-HSL in yeast by BjaI, MesI and RpaI.** LC-MS chromatogram of a standard of C12-HSL, supernatant of *S. cerevisiae* strains ACA046 (BjaI), ACA050 (RpaI) and ACA055 (MesI), and of a negative control. Yeast strains were grown for 24 h on fed batch medium. Supernatant was extracted using ethyl acetate and dried extracts were dissolved in methanol before analysis. C12-HSL eluted at a retention time of 3.6 min.

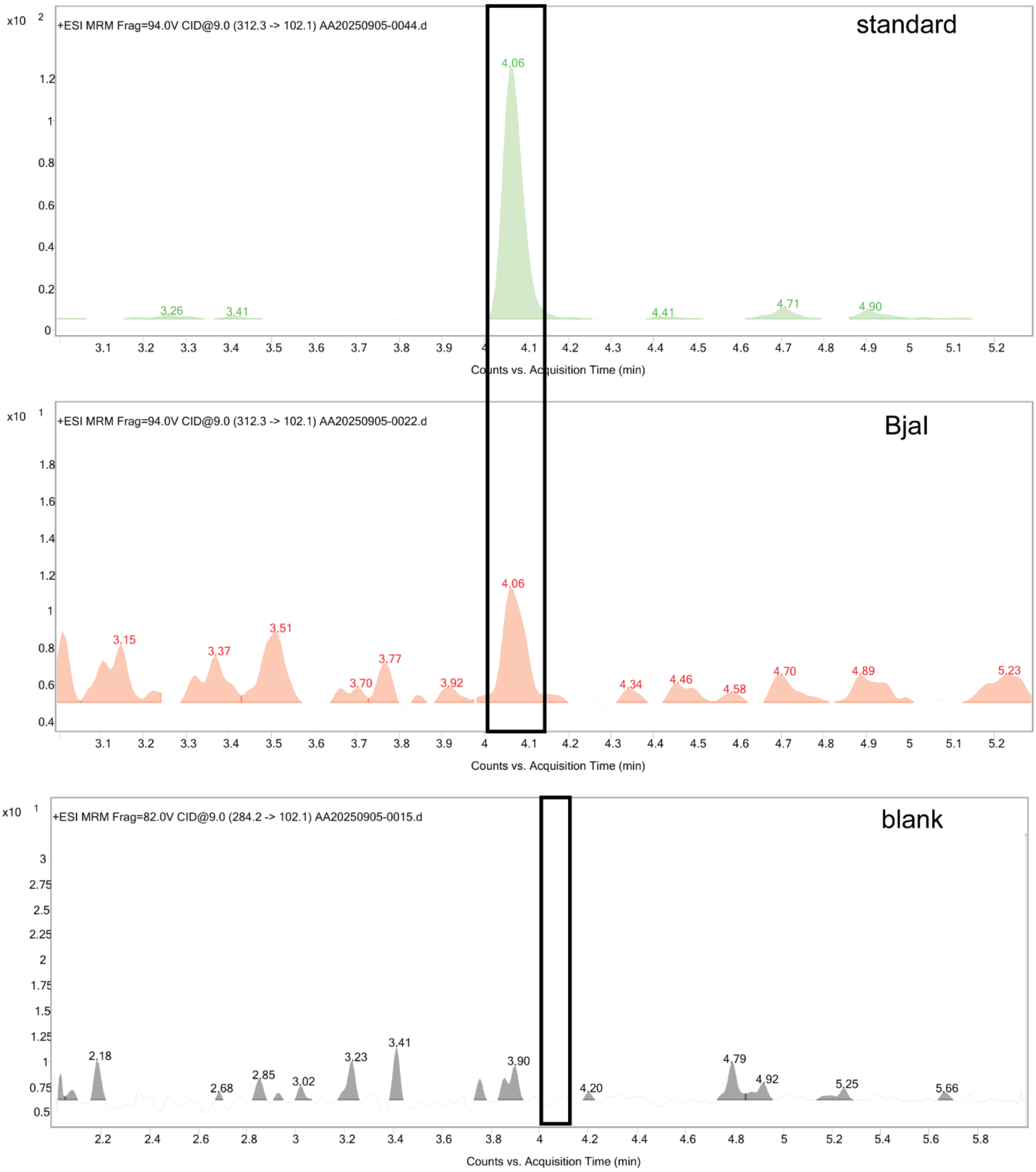

**Figure S6: Production of C14-HSL in yeast by *BjaI*.** LC-MS chromatogram of a standard of C14-HSL, supernatant of *S. cerevisiae* strain ACA046 (*BjaI*) and of a negative control. Yeast strains were grown for 24 h on fed batch medium. Supernatant was extracted using ethyl acetate and dried extracts were dissolved in methanol before analysis. C14-HSL eluted at a retention time of 4.06 min.

**Table S1: Overview of AHLs produced by different synthases introduced in yeast.** Strains ACA049 (*EsaI*), ACA046 (*BjaI*), ACA071 (*BjaI*-PRX), ACA047 (BraI) ACA072 (BraI-PRX), ACA050 (*RpaI*), ACA070 (*RpaI*-PRX), ACA055 (*MesI*), ACA066 (*MesI*-PRX) and ACA056 (*MplI*) were grown in duplicate for 24 h on fedbatch medium and supernatant was analysed by LC-MS to determine the AHL production. Mean and standard deviation of the mean are shown.

| **Synthase** | **C4-HSL** | **C6-HSL** | **C8-HSL** | **C10-HSL** |
|---|---|---|---|---|
| **EsaI** | - | - | - | - |
| **BjaI** | 2468 ± 112 nM | 37 ± 2 nM | 4 ± 0 nM | 11 ± 0 nM |
| **BjaI-PRX** | 13 ± 0 nM | 52 ± 2 nM | 8 ± 0 nM | 146 ± 2 nM |
| **BraI** | - | - | - | - |
| **BraI-PRX** | - | - | - | - |
| **RpaI** | - | 3 ± 1 nM | 6 ± 1 nM | - |
| **RpaI-PRX** | - | 2 ± 0 nM | 4 ± 0 nM | 6 ± 1 nM |
| **MesI** | 1 ± 0 nM | 41 ± 9 nM | 2 ± 0 nM | 3 ± 2 nM |
| **MesI-PRX** | 2 ± 1 nM | 173 ± 7 nM | 56 ± 1 nM | 229 ± 17 nM |
| **MplI** | - | - | - | - |

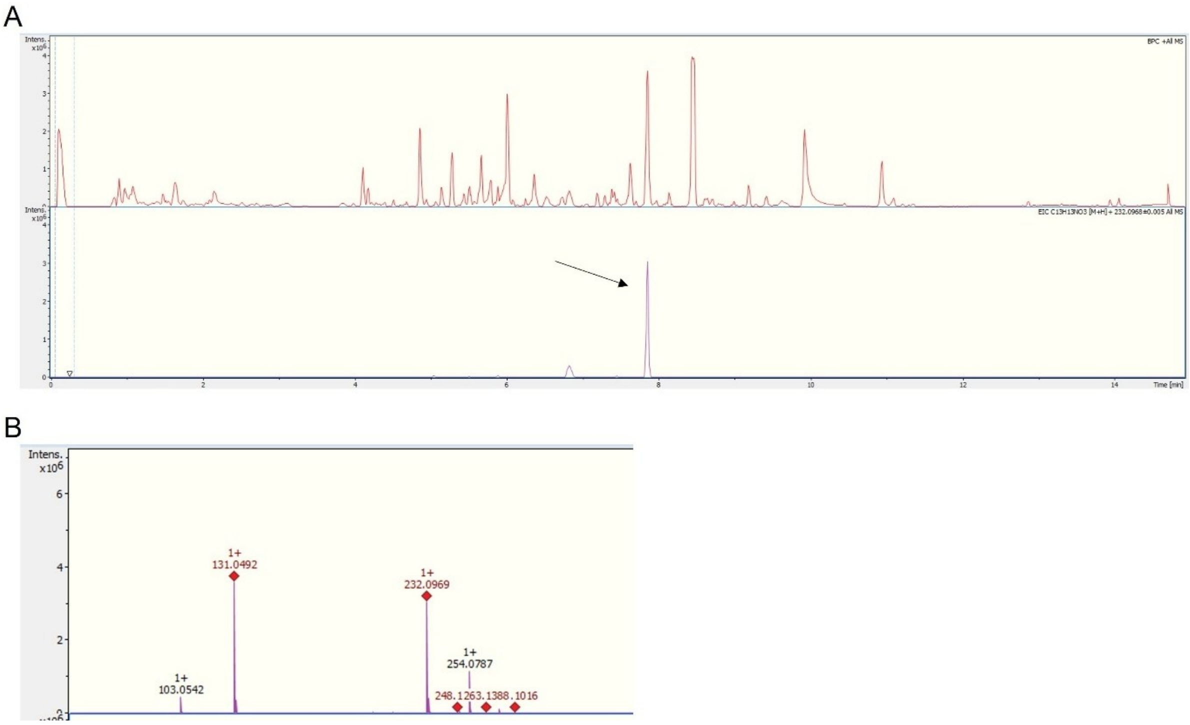

**Figure S7: Cinnamoyl-HSL peak and MS2 fragmentation pattern. (A)** Yeast strains AAA119 and AAA121 were grown for 24 h on fed batch medium**.** Supernatant was analysed by LC-MS and LC-MS chromatogram of one representative sample is shown. **(B)** MS/MS fragmentation spectra of cinnamoyl-HSL (*m/z* 232) acquired at 7.9 min. Major fragments include *m/z* 131 and *m/z* 103 [3].

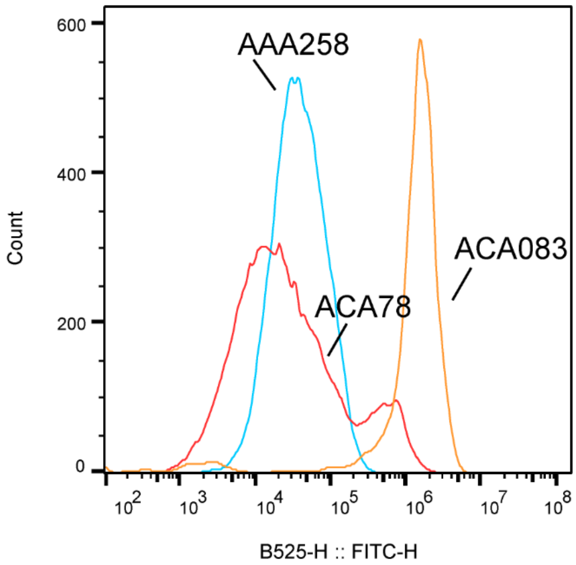

**Figure S8: Population fluorescence of VP48-*rpaR* library as determined by flow cytometry.** Using ep-PCR, a library was created of VP48-*rpaR* and this library was integrated into a yeast strain containing *amdS* and yeGFP under control of p*GAL1*_core_5x*rpaO*. The yeast library was grown for 2 days on transformation selection media (SC -HIS +NAT) and transferred one more time for overnight growth on fresh transformation selection medium. Next, the population of selected transformants was transferred to counter-selection media (SMD + F-Ac) supplemented with 400 nM C10-HSL and grown for 1 day. Subsequently, cells were transferred twice on selection media (SMD -N + acetamide), containing 1 µM 3-*p*-coumaroyl-HSL. This final enriched population (ACA78) together with the parental biosensor strain (AAA258) and a selected colony from the library population (ACA083), were grown in the presence of 1 µM 3-oxo-C8-HSL and analysed after 6 h on the flow cytometer.

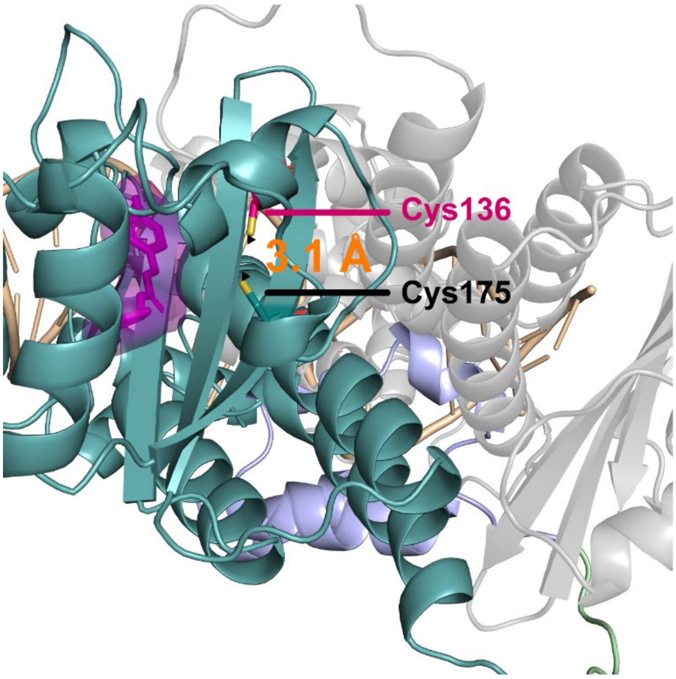

**Figure S9: Structural analysis of RpaR*.** Close-up view of the RpaR* G88C substitution showing spatial proximity to Cys175 compatible with possible disulfide bond formation.

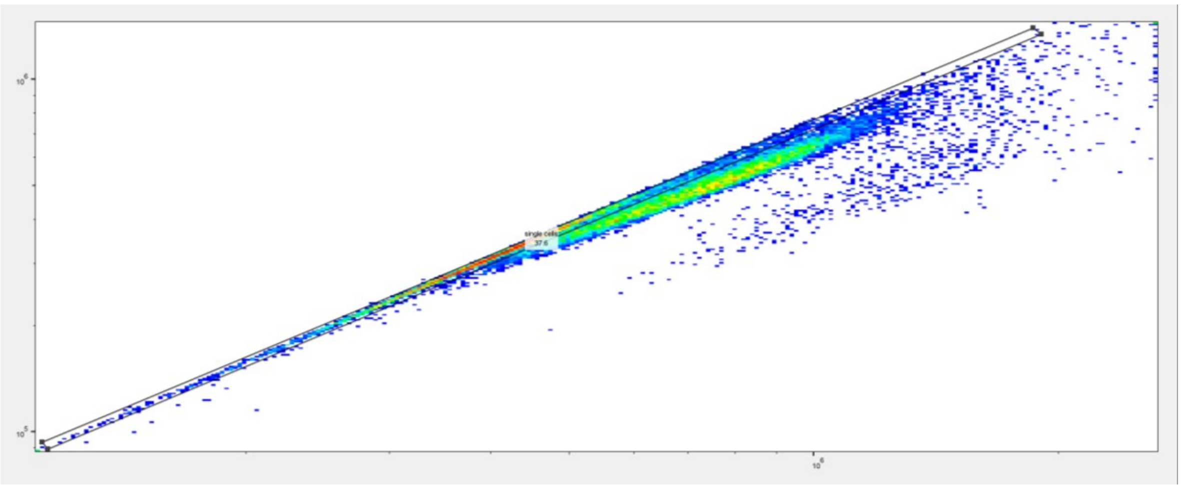

**Figure S10: Example of gating strategy for flow cytometry data.** FSC-A was plotted against FSC-H to gate for singlets events.

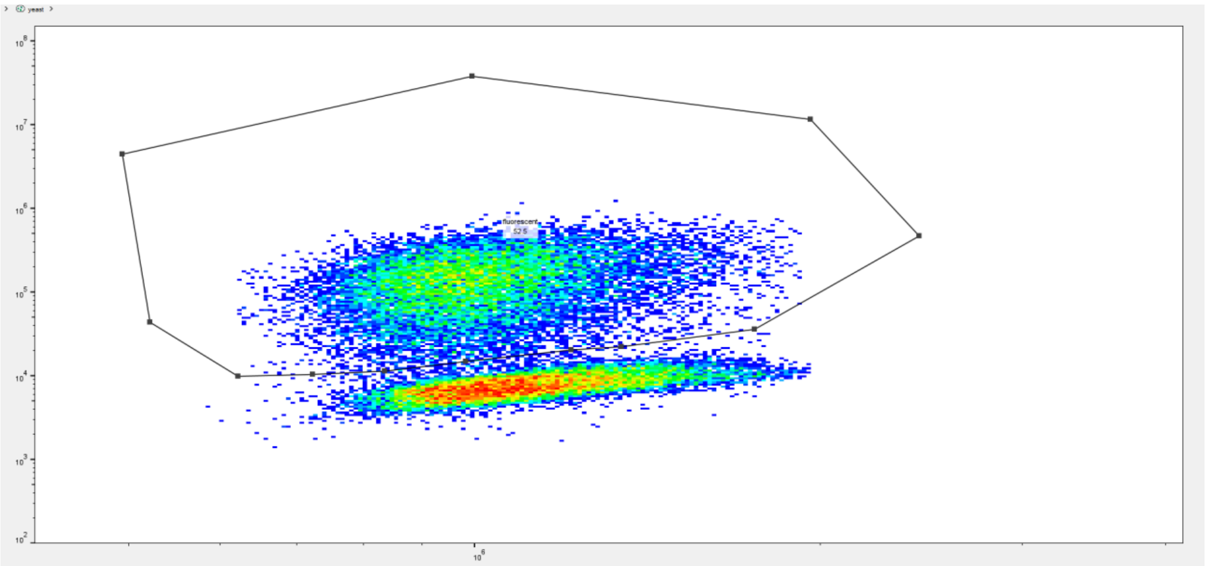

**Figure S11: Example of gating strategy for flow cytometry data for co-culture.** After gating for singlets events (Figure S10), either the red or the green channel was plotted against FSC-H and a gate was drawn around the fluorescent subpopulation to determine the median fluorescence of each specific (green and red) subpopulation.

**Table S2: overview of yeast strains used in this study and details about the strain construction**

| Name | Relevant genotype | Details strain construction | source |
|---|---|---|---|
| AAA001 | *MAT*-a *ura3 his3 leu2 trp1 p-Cas9* | - | [2] |
| AAA006 | *MAT*-a *ura3 his3 leu2 trp1* XI-3-p*TEF1*-*esaO*_105-yeGFP-t*CPS1*, *p-Cas9* | AAA001: pCfb6915 + NotI digested pAvA015 | This study |
| AAA010 | *MAT*-a *ura3 his3 leu2 trp1* XI-3-*pGAL1*-5x*luxO*-yeGFP-t*CPS1*, *p-Cas9* | - | [2] |
| AAA015 | *MAT*-a *ura3 his3 leu2 trp1* XI-3-p*TEF1*-*esaO*_105-yeGFP-t*CPS1* | AAA006: plasmids removed | This study |
| AAA019 | *MAT*-a *ura3 his3 leu2 trp1* XI-3-*pGAL1*-5x*luxO*-yeGFP-t*CPS1* | - | [2] |
| AAA031 | *MAT*-a *ura3 his3 leu2 trp1* XI-3-p*TEF1*-*esaO*_105-yeGFP-t*CPS1*, X-4-p*PGK1*-*esaR*$_{V220A}$-t*ADH1* | AAA006: pCfB3042 + NotI digested pAvA039 | This study |
| AAA032 | *MAT*-a *ura3 his3 leu2 trp1* XI-3-p*TEF1*-*esaO*_105-yeGFP-t*CPS1*, X-4-p*PGK1*-*esaR*$_{D91G}$-t*ADH1* | AAA006: pCfB3042+ NotI digested pAvA040 | This study |
| AAA033 | *MAT*-a *ura3 his3 leu2 trp1* XI-3-p*TEF1*-*esaO*_105-yeGFP-t*CPS1*, X-4-p*PGK1*-*esaR*-t*ADH1* | AAA006: pCfB3042 + NotI digested pAvA041 | This study |
| AAA084 | *MAT*-a *ura3 his3 leu2 trp1* X-2-p*TDH3*-*rpaI*_co-t*CYC1*, *p-Cas9* | AAA001: pCfB3020 + NotI digested pAvA081 | This study |
| AAA085 | *MAT*-a *ura3 his3 leu2 trp1* X-2-p*TDH3*-*rpaI*_co-t*CYC1* | AAA084: plasmids removed | This study |
| AAA089 | *MAT*-a *ura3 his3 leu2 trp1* XI-1-t*ADH1*-At4CL1-p*ACT1*--p*PGI1*-FjTAL-t*CYC1*, pCas9 | AAA001: pCfB3043 + NotI digested pAvA070 | This study |
| AAA092 | *MAT*-a *ura3 his3 leu2 trp1* X-2-p*TDH3*-*rpaI*_co-t*CYC1*, XI-1-t*ADH1*-At4CL1-p*ACT1*--p*PGI1*-FjTAL-t*CYC1* | AAA084: pCfB3043 + NotI digested pAvA070 | This study |
| AAA095 | *MAT*-a *ura3 his3 leu2 trp1* XI-3-p*TEF1*_*luxO*-yeGFP-t*CPS1*, *p-Cas9* | - | [2] |
| AAA119 | *MAT*-a *ura3 his3 leu2 trp1* X-2-p*TDH3*-*rpaI*_co-t*CYC1*, XI-1-t*ADH1*-At4CL1-p*ACT1*--p*PGI1*-AtPAL-t*CYC* | AAA084: pCfB3043 + NotI digested pAvA122 | This study |
| AAA121 | *MAT*-a *ura3 his3 leu2 trp1* X-2-p*TDH3*-*rpaI*_co-t*CYC1*, XI-1-t*ADH1*-At4CL2-p*ACT1*--p*PGI1*-AtPAL-t*CYC* | AAA084: pCfB3043 + NotI digested pAvA123 | This study |
| AAA122 | *MAT*-a *ura3 his3 leu2 trp1* X-2-p*TDH3*-*rpaI*_co-t*CYC1*, XI-1-t*ADH1*-At4CL2-p*ACT1*--p*PGI1*-FjTAL-t*CYC*, *p-Cas9* | AAA084: pCfB3043 + NotI digested pAvA124 | This study |
| AAA123 | *MAT*-a *ura3 his3 leu2 trp1* X-2-p*TDH3*-*rpaI*_co-t*CYC1*, XI-1-t*ADH1*-At4CL2-p*ACT1*--p*PGI1*-FjTAL-t*CYC* | AAA122: plasmids removed | This study |
| AAA144 | *MAT*-a *ura3 his3 leu2 trp1* XI-3-p*TEF1*_2x*esaO*_98_105-yeGFP-t*CPS1*, *p-Cas9* | AAA001: pCfB3045 + NotI digested pAvA133 | This study |
| AAA145 | *MAT*-a *ura3 his3 leu2 trp1* XI-3-p*TEF1*_2x*esaO*_98_105-yeGFP-t*CPS1* | AAA144: plasmids removed | This study |
| AAA156 | *MAT*-a *ura3 his3 leu2 trp1* XI-3-*pGAL1*-5x*luxO*-yeGFP-t*CPS1*, X-4-p*PGK1*-*GAL4*$_{AD}$-*luxR*_gen1_t*ADH1* | - | [2] |
| AAA167 | *MAT*-a *ura3 his3 leu2 trp1* XI-3-p*TEF1*_2x*esaO*_98_105-yeGFP-t*CPS1*, X-4-p*PGK1*-*esaR*$_{D91G}$-t*ADH1* | AAA144: pCfB3042 + NotI digested pAvA040 | This study |

| | | | |
|---|---|---|---|
| AAA159 | *MAT*-a *ura3 his3 leu2 trp1* XI-3-*pGAL1*-5x*rpaO*-yeGFP-t*CPS1*, pCas9 | AAA001: pCfB3045 + NotI digested pAvA145 | This study |
| AAA160 | *MAT*-a *ura3 his3 leu2 trp1* XI-3-*pGAL1*-5x*rpaO*-yeGFP-t*CPS1* | AAA159: plasmids removed | This study |
| AAA212 | *MAT*-a *ura3 his3 leu2 trp1* XI-3-p*CCW-esaO*-yeGFP-t*CPS1*, *p-Cas9* | AAA001: pCfB3045 + NotI digested pAvA160 | This study |
| AAA213 | *MAT*-a *ura3 his3 leu2 trp1* XI-3-p*CCW-esaO*-yeGFP-t*CPS1* | AAA212: plasmids removed | This study |
| AAA214 | *MAT*-a *ura3 his3 leu2 trp1* X-2-p*TDH3-bjaI*_co-t*CYC1* | AAA001: pCfB3020 + NotI digested pAvA158 | This study |
| AAA215 | *MAT*-a *ura3 his3 leu2 trp1* X-2-p*TDH3-braI*_co-t*CYC1* | AAA001: pCfB3020 + NotI digested pAvA159 | This study |
| AAA216 | *MAT*-a *ura3 his3 leu2 trp1* XI-3-p*CCW-esaO*-yeGFP-t*CPS1*, X-4-p*PGK1-esaR*$_{D91G}$-t*ADH1* | AAA212: pCfB3042 + NotI digested pAvA040 | This study |
| AAA218 | *MAT*-a *ura3 his3 leu2 trp1* XI-3-p*CCW-esaO*-yeGFP-t*CPS1*, X-4-p*PGK1-esaR*$_{D91G}$-NLS-t*ADH1* | AAA212: pCfB3042 + NotI digested pAvA157 | This study |
| AAA236 | *MAT*-a *ura3 his3 leu2 trp1* X-2-p*TDH3-rpaI*_co-t*CYC1*, XI-1-t*ADH1*-At4CL2-p*ACT1*--p*PGI1*-FjTAL-t*CYC*, XI-3-*pGAL1*-5x*rpaO*-yeGFP-t*CPS1*, *p-Cas9* | AAA122: pCfB3045 + NotI digested pAvA145 | This study |
| AAA237 | *MAT*-a *ura3 his3 leu2 trp1* X-2-p*TDH3-rpaI*_co-t*CYC1*, XI-1-t*ADH1*-At4CL2-p*ACT1*--p*PGI1*-FjTAL-t*CYC*, XI-3-*pGAL1*-5x*rpaO*-yeGFP-t*CPS1* | AAA236: plasmids removed | This study |
| AAA250 | *MAT*-a *ura3 his3 leu2 trp1* X-2-p*TDH3-rpaI*_co-t*CYC1*, XI-1-t*ADH1*-At4CL2-p*ACT1*--p*PGI1*-FjTAL-t*CYC*, XI-3-*pGAL1*-5x*luxO*-yeGFP-t*CPS1*, X-4-p*PGK1*- *GAL4*$_{AD}$ -*luxR*$_{N86K}$-t*ADH1* | AAA234: pCfB3042 + X-4-p*PGK1*-GAL4_AD-NLS-*luxR*_N86K + *luxR*_N86K-t*ADH1* | This study |
| AAA257 | *MAT*-a *ura3 his3 leu2 trp1* XI-3-*pGAL1*-5x*rpaO*-yeGFP-t*CPS1*, X-4-p*PGK1*-VP48-*rpaR*-t*ADH1*, pCas9 | AAA159: pCfB3042 + NotI digested pAvA169 | This study |
| AAA258 | *MAT*-a *ura3 his3 leu2 trp1* XI-3-*pGAL1*-5x*rpaO*-yeGFP-t*CPS1*, X-4-p*PGK1*-VP48-*rpaR*-t*ADH1* | AAA257: plasmids removed | This study |
| AAA261 | *MAT*-a *ura3 his3 leu2 trp1* X-2-p*TDH3-rpaI*_co-t*CYC1*, XI-1-t*ADH1*-At4CL2-p*ACT1*--p*PGI1*-FjTAL-t*CYC*, XI-3-*pGAL1*-5x*luxO*-yeGFP-t*CPS1*, X-4-p*PGK1*- *GAL4*$_{AD}$ -*luxR*_gen1_t*ADH1* | AAA234: pCfB3042 + NotI digested pAvA139 | This study |
| AAA263 | *MAT*-a *ura3 his3 leu2 trp1* X-2-p*TDH3-rpaI*_co-t*CYC1*, XI-1-t*ADH1*-At4CL2-p*ACT1*--p*PGI1*-FjTAL-t*CYC*, XI-3-*pGAL1*-5x*rpaO*-yeGFP-t*CPS1*, X-4-p*PGK1*-VP48-*rpaR*-t*ADH1* | AAA236: pCfB3042 + NotI digested pAvA169 | This study |
| AAA264 | *MAT*-a *ura3 his3 leu2 trp1* X-2-p*TDH3-bjaI*_co-t*CYC1*, XI-3-*pGAL1*-5x*rpaO*-yeGFP-t*CPS1*, X-4-p*PGK1*-VP48-*rpaR*-t*ADH1* | AAA240: pCfB3042 + NotI digested pAvA169 | This study |
| AAA268 | *MAT*-a *ura3 his3 leu2 trp1* XI-3-*pGAL1*-5x*lasO*-yeGFP-t*CPS1*, *p-Cas9* | AAA001: pCfB3045 + NotI digested pAvA187 | This study |
| AAA269 | *MAT*-a *ura3 his3 leu2 trp1* XI-3-*pGAL1*-5x*lasO*-yeGFP-t*CPS1* | AAA268: plasmids removed | This study |
| AAA270 | *MAT*-a *ura3 his3 leu2 trp1* XI-3-*pGAL1*-5x*traO*-yeGFP-t*CPS1*, *p-Cas9* | AAA001: pCfB3045 + NotI digested pAvA185 | This study |
| AAA271 | *MAT*-a *ura3 his3 leu2 trp1* XI-3-*pGAL1*-5x*traO*-yeGFP-t*CPS1* | AAA270: plasmids removed | This study |
| AAA274 | *MAT*-a *ura3 his3 leu2 trp1* XI-3-*pGAL1*-5x*lasO*-yeGFP-t*CPS1*, X-4-p*PGK1*-VP48-*lasR*-t*ADH1* | AAA268: pCfB3042 + NotI digested pAvA173 | This study |
| AAA275 | *MAT*-a *ura3 his3 leu2 trp1* XI-3-*pGAL1*-5x*traO*-yeGFP-t*CPS1*, X-4-p*PGK1*-VP48-*traR*-t*ADH1* | AAA270: pCfB3042 + NotI digested pAvA171 | This study |

| | | | |
|---|---|---|---|
| AAA282 | MAT-a ura3 his3 leu2 trp1 XI-3- pTEF1-luxO_105-yeGFP-tCPS1, X-4-pPGK1- $GAL4_{AD}$ -luxR_gen1_tADH1, p-Cas9 | AAA095: pCfB3042 + NotI digested pAvA139 | This study |
| AAA289 | MAT-a ura3 his3 leu2 trp1 XI-3- pTEF1-luxO_105-yeGFP-tCPS1, X-4-pPGK1- $GAL4_{AD}$ -luxR_gen1_tADH1, XII-5-pGAL1_5xluxO-mKate2-tCPS1 | AAA282: pCfB3050 + NotI digested pAvA197 | This study |
| AAA295 | MAT-a ura3 his3 leu2 trp1 XI-3-pTEF1-luxO_105-yeGFP-tCPS1, XII-5-pGAL1_5xluxO-mKate2-tCPS1 | AAA095: pCfB3050 + NotI digested pAvA197 | This study |
| AAA296 | MAT-a ura3 his3 leu2 trp1 XI-3-pGAL1-5xluxO-yeGFP-tCPS1, X-2-pTDH3-mesI-PRX | AAA010: pCfB3020 + NotI digested pAvA199 | This study |
| AAA298 | MAT-a ura3 his3 leu2 trp1 XI-3-pGAL1-5xrpaO-yeGFP-tCPS1, X-2-pTDH3-mesI-PRX | AAA159: pCfB3020 + NotI digested pAvA199 | This study |
| AAA300 | MAT-a ura3 his3 leu2 trp1 XI-3-pGAL1-5xrpaO-yeGFP-tCPS1, X-4-pPGK1-VP48-rpaR-tADH1, X-2-pTDH3-mesI-PRX | AAA257: pCfB3020 + NotI digested pAvA199 | This study |
| AAA309 | MAT-a ura3 his3 leu2 trp1 X-2-pTDH3-bjaI_co-tCYC, XII-5-tADH1-SAM2-pTEF1--pPGK1-MET6-tCYC1 | AAA214: p-Cas9 + pCfB3050 + NotI digested pAvA118 | This study |
| AAA310 | MAT-a ura3 his3 leu2 trp1 X-2-pTDH3-mesI-PRX-tCYC, XII-5-tADH1-SAM2-pTEF1--pPGK1-MET6-tCYC1, p-Cas9 | AAA001: pCfB3020 + NotI digested pAvA199 + pCfB3050 + NotI digested pAvA118 | This study |
| AAA311 | MAT-a ura3 his3 leu2 trp1 X-2-pTDH3-mesI-PRX-tCYC, XII-5-tADH1-SAM2-pTEF1--pPGK1-MET6-tCYC1 | AAA310: plasmids removed | This study |
| AAA312 | MAT-a ura3 his3 leu2 trp1 X-2-pTDH3-rpaI_co-tCYC1, XI-1-tADH1-At4CL2-pACT1--pPGI1-FjTAL-tCYC, XI-3-pGAL1-5xluxO-mKate2-tCPS1, X-4-pPGK1- $GAL4_{AD}$ -$luxR_{N86K}$-tADH1 | AAA122: pCfB6915 + NotI digested pAvA179 + pCfB3042 + X-4-pPGK1-GAL4_AD-NLS-luxR_N86K + luxR_N86K-tADH1-X-4 | This study |
| AAA313 | MAT-a ura3 his3 leu2 trp1 X-2-pTDH3-rpaI_co-tCYC1, XI-1-tADH1-At4CL2-pTEF1--pPGK1-FjTAL-tCYC1, p-Cas9 | AAA084: pCfB3043 + NotI digested pAvA205 | This study |
| AAA314 | MAT-a ura3 his3 leu2 trp1 X-2-pTDH3-rpaI_co-tCYC1, XI-1-tADH1-At4CL2-pTEF1--pPGK1-FjTAL-tCYC1 | AAA313: plasmids removed | This study |
| AAA319 | MAT-a ura3 his3 leu2 trp1 X-2-pTDH3-RpaI_co-tCYC1, XI-1-tADH1-At4CL2-pACT1--pPGI1-FjTAL-tCYC, XI-3-pGAL1-5xluxO-mKate2-tCPS1, X-4-pPGK1-$GAL4_{AD}$ -luxR_gen1_tADH1 | AAA122: pCfB6915 + NotI digested pAvA179, pCfB3042 + NotI digested pAvA139 | This study |
| AAA320 | MAT-a ura3 his3 leu2 trp1 X-2-pTDH3-mesI-PRX-tCYC, XII-5-tADH1-SAM2-pTEF1--pPGK1-MET6-tCYC1, XI-3-pGAL1-5xrpaO-yeGFP-tCPS1, X-4-pPGK1-VP48-rpaR-tADH1 | AAA310: pCfB6915 + NotI digested pAvA145, pCfB3042 + NotI digested pAvA169 | This study |
| AAA321 | MAT-a ura3 his3 leu2 trp1 XI-3-pGAL1-5xrpaO-eGFP-tCPS1, X-4-pPGK1-VP48-rpaR*-tADH1 | AAA159: pCfB3042 + X-4-pPGK1-VP48-rpaR*-tADH1-X-4 | This study |
| AAA322 | MAT-a ura3 his3 leu2 trp1 X-2-pTDH3-rpaI_co-tCYC1, XI-1-tADH1-At4CL2-pTEF1--pPGK1-FjTAL-tCYC, XI-3-pGAL1-5xluxO-mKate2-tCPS1, X-4-pPGK1-$GAL4_{AD}$ -luxR_gen1_tADH1 | AAA313: pCfB6915 + NotI digested pAvA179, pCfB3042 + NotI digested pAvA139 | This study |
| AAA323 | MAT-a ura3 his3 leu2 trp1 X-2-pTDH3-mesI-PRX-tCYC, XII-5-tADH1-SAM2-pTEF1--pPGK1-MET6-tCYC1, XI-3-pGAL1-5xrpaO-yeGFP-tCPS1, X-4-pPGK1-VP48-rpaR* -tADH1 | AAA310: pCfB6915 + NotI digested pAvA145, pCfB3042 + X-4-pPGK1-VP48-rpaR*-tADH1-X-4 | This study |
| ACA001 | MAT-α URA3 his3 LEU2 TRP1 p-Cas9 | - | [2] |

| | | | |
|---|---|---|---|
| ACA040 | MAT-α URA3 his3 LEU2 TRP1 XI-3-pGAL1-5xrpaO-yeGFP-tCPS1, p-Cas9 | ACA001: pCfB3045 + NotI digested pAvA145 | This study |
| ACA042 | MAT-α URA3 his3 LEU2 TRP1 XI-3-pGAL1-5xrpaO-yeGFP-tCPS1, XII-5-pGAL1-5xrpaO-amdS-tCPS1, p-Cas9 | ACA040: pCfB3050 + NotI digested pAvA146 | This study |
| ACA015 | MAT-α URA3 his3 LEU2 TRP1 XI-3-pGAL1_core-5xluxO-amdS-tCPS1 XII-5-pGAL1_core-5xluxO-yeGFP-tCPS1 p-Cas9 | - | [2] |
| ACA044 | MAT-α URA3 his3 LEU2 TRP1 XI-3-pGAL-5xluxO-amdS-tCPS1, XII-5-pGAL1-5xluxO-yeGFP-tCPS1, VIII-1-tADH1-SAM2-pTEF1--pPGK1-MET6-tCYC1, p-Cas9 | ACA015: pCfB9340 + NotI digested pAvA155 | This study |
| ACA045 | MAT-α URA3 his3 LEU2 TRP1 XI-3-pGAL-5xluxO-amdS-tCPS1, XII-5-pGAL1-5xluxO-yeGFP-tCPS1, VIII-1-tADH1-SAM2-pTEF1--pPGK1-MET6-tCYC1, X-4-pPGK1- GAL4$_{AD}$ -luxR_gen1-tADH1, p-Cas9 | ACA044: pCfB3042 + NotI digested pAvA139 | This study |
| ACA046 | MAT-α URA3 his3 LEU2 TRP1 XI-3-pGAL1-5xluxO-amdS-tCPS1, XII-5-pGAL1-5xluxO-yeGFP-tCPS1, VIII-1-tADH1-SAM2-pTEF1--pPGK1-MET6-tCYC1, X-4-pPGK1- GAL4$_{AD}$ -luxR_gen1-tADH1, X-2-pTDH3-bjaI-tCYC, p-Cas9 | ACA045: pCfB3020 + NotI digested pAvA158 | This study |
| ACA047 | MAT-α URA3 his3 LEU2 TRP1 XI-3-pGAL-5xluxO-amdS-tCPS1, XII-5-pGAL1-5xluxO-yeGFP-tCPS1, VIII-1-tADH1-SAM2-pTEF1--pPGK1-MET6-tCYC1, X-4-pPGK1- GAL4$_{AD}$ -luxR_gen1-tADH1, X-2-pTDH3-BraI-tCYC, p-Cas9 | ACA045: pCfB3020 + NotI digested pAvA159 | This study |
| ACA049 | MAT-α URA3 his3 LEU2 TRP1 XI-3-pGAL-5xluxO-amdS-tCPS1, XII-5-pGAL1-5xluxO-yeGFP-tCPS1, VIII-1-tADH1-SAM2-pTEF1--pPGK1-MET6-tCYC1, X-4-pPGK1- GAL4$_{AD}$ -luxR_gen1-tADH1, X-2-pTDH3-esaI-tCYC, p-Cas9 | ACA045: pCfB3020 + NotI digested pAvA079 | This study |
| ACA050 | MAT-α URA3 his3 LEU2 TRP1 XI-3-pGAL-5xluxO-amdS-tCPS1, XII-5-pGAL1-5xluxO-yeGFP-tCPS1, VIII-1-tADH1-SAM2-pTEF1--pPGK1-MET6-tCYC1, X-4-pPGK1- GAL4$_{AD}$ -luxR_gen1-tADH1, X-2-pTDH3-rpaI-tCYC, p-Cas9 | ACA045: pCfB3020 + NotI digested pAvA081 | This study |
| ACA055 | MAT-α URA3 his3 LEU2 TRP1 XI-3-pGAL-5xluxO-amdS-tCPS1, XII-5-pGAL1-5xluxO-yeGFP-tCPS1, VIII-1-tADH1-SAM2-pTEF1--pPGK1-MET6-tCYC1, X-4-pPGK1- GAL4$_{AD}$ -luxR_gen1-tADH1, X-2-pTDH3-mesI-tCPS1, p-Cas9 | ACA045: pCfB3020 + NotI digested pAvA175 | This study |
| ACA056 | MAT-α URA3 his3 LEU2 TRP1 XI-3-pGAL1-5xluxO-amdS-tCPS1, XII-5-pGAL1-5xluxO-yeGFP-tCPS1, VIII-1-tADH1-SAM2-pTEF1--pPGK1-MET6-tCYC1, X-4-pPGK1- GAL4$_{AD}$ -luxR_gen1-tADH1, X-2-pTDH3-mplI-tCPS1, p-Cas9 | ACA045: pCfB3020 + NotI digested pAvA176 | This study |
| ACA065 | MAT-α URA3 his3 LEU2 TRP1 XI-3-pGAL1-5xtraO-yeGFP-tCPS1, p-Cas9 | ACA001: pCfB3045 + NotI digested pAvA195 | This study |

| | | | |
|---|---|---|---|
| ACA066 | MAT-α URA3 his3 LEU2 TRP1 XI-3-pGAL1-5xluxO-amdS-tCPS1, XII-5-pGAL1-5xluxO-yeGFP-tCPS1, VIII-1-tADH1-SAM2-pTEF1--pPGK1-MET6-tCYC1, X-4-pPGK1- $GAL4_{AD}$ -luxR_gen1-tADH1, X-2-pTDH3-mesI-PRX-tCPS1, p-Cas9 | ACA045: pCfB3020 + NotI digested pAvA199 | This study |
| ACA069 | MAT-α URA3 his3 LEU2 TRP1 XI-3-pGAL1-5xtraO-yeGFP-tCPS1, XII-5-pGAL1-5xtraO-amdS-tCPS1, p-Cas9 | ACA065: pCfB3050 + NotI digested pAvA195 | This study |
| ACA070 | MAT-α URA3 his3 LEU2 TRP1 XI-3-pGAL1-5xluxO-amdS-tCPS1, XII-5-pGAL1-5xluxO-yeGFP-tCPS1, VIII-1-tADH1-SAM2-pTEF1--pPGK1-MET6-tCYC1, X-4-pPGK1- $GAL4_{AD}$ -luxR_gen1-tADH1, X-2-pTDH3-rpaI-PRX-tCPS1, p-Cas9 | ACA045: pCfB3020 + NotI digested pAvA202 | This study |
| ACA071 | MAT-α URA3 his3 LEU2 TRP1 XI-3-pGAL1-5xluxO-amdS-tCPS1, XII-5-pGAL1-5xluxO-yeGFP-tCPS1, VIII-1-tADH1-SAM2-pTEF1--pPGK1-MET6-tCYC1, X-4-pPGK1- $GAL4_{AD}$ -luxR_gen1-tADH1, X-2-pTDH3-bjaI-PRX-tCPS1, p-Cas9 | ACA045: pCfB3020 + NotI digested pAvA200 | This study |
| ACA072 | MAT-α URA3 his3 LEU2 TRP1 XI-3-pGAL1-5xluxO-amdS-tCPS1, XII-5-pGAL1-5xluxO-yeGFP-tCPS1, VIII-1-tADH1-SAM2-pTEF1--pPGK1-MET6-tCYC1, X-4-pPGK1- $GAL4_{AD}$ -luxR_gen1-tADH1, X-2-pTDH3-braI-PRX-tCPS1, p-Cas9 | ACA045: pCfB3020 + NotI digested pAvA201 | This study |
| ACA074 | MAT-α URA3 his3 LEU2 TRP1 XI-3-pGAL1-5xtraO-eGFP-tCPS1, XII-5-pGAL1-5xtraO-amdS-tCPS1, X-4-pPGK1-VP48-traR-tADH1 (library), p-Cas9 | ACA069: pCfB3042 + X-4-pPGK1 + VP48-traR (library) + tADH1-X-4 | This study |
| ACA078 | MAT-α URA3 his3 LEU2 TRP1 XI-3-pGAL1-5xrpaO-eGFP-tCPS1, XII-5-pGAL1-5xrpaO-amdS-tCPS1, X-4-pPGK1-VP48-rpaR-tADH1 (library), p-Cas9 | ACA042: pCfB3042 + X-4-pPGK1 + VP48-traR (library) + tADH1-X-4 | This study |
| ACA080 | MAT-α URA3 his3 LEU2 TRP1 XI-3-pGAL1-5xtraO-eGFP-tCPS1, XII-5-pGAL1-5xtraO-AmdS-tCPS1, X-4-pPGK1-VP48-traR-tADH1_colony 2 | Mutant selected from ACA074 | This study |
| ACA083 | MAT-α URA3 his3 LEU2 TRP1 XI-3-pGAL1-5xrpaO-eGFP-tCPS1, XII-5-pGAL1-5xrpaO-AmdS-tCPS1, X-4-pPGK1-VP48-rpaR-tADH1_colony D10 | Mutant selected from ACA078 | This study |

**Table S3: plasmids used in this study**

| Name | Details | Origin |
|---|---|---|
| pTAJAK-161 | Cas9, HIS | [4] |
| pCfB3045 | XI-3 gRNA, NAT | [5] |
| pCfB6915 | XI-3 gRNA, URA | [6] |
| pCfB3042 | X-4 gRNA, NAT | [5] |
| pCfB6912 | X-4 gRNA, URA | [6] |
| pCfB9340 | VIII-1 gRNA, NAT | [7] |
| pCfB3050 | XII-5 gRNA, NAT | [5] |
| pCfB3020 | X-2 gRNA, NAT | [5] |
| pCfB6910 | X-2 gRNA, URA | [6] |
| pCfB2904 | XI-3 USER overhangs | [5] |
| pCfB3035 | X-4 USER overhangs | [5] |
| pCfB2909 | XII-5 USER overhangs | [5] |
| pCfB2899 | X-2 USER overhangs | [5] |
| pCfB9359 | VIII-1 USER overhangs | [7] |
| pCfB756 | FjPAL | [8] |
| pCfB757 | At4CL1 | [8] |
| pCfB758 | At4CL2 | [8] |
| pCfB873 | FjTAL | [9] |
| pAC-*EsaR-EsaI* | *esaR* | [10] |
| pAC-*EsaR*-V220A | *esaR* | [10] |
| pAC-*EsaR*-D91G | *esaR* | [10] |
| Bsrs078-*LasR* | *lasR* | [11] |
| Bsrs078-*TraR* | *traR* | [11] |
| Bsrs103-*RpaR-RpaI* | *rpaR* | [11] |
| pAvA015 | XI-3-p*TEF1*-*esaO*1_105-yeGFP-t*CPS1* | This study |
| pAvA026 | XI-3-p*GAL1*-5x*luxO*-yeGFP-tCPS1 | [2] |
| pAvA039 | X-4-p*PGK1*-*esaR*-V220A-t*ADH1* | This study |
| pAvA040 | X-4-p*PGK1*-*esaR*-D91G-t*ADH1* | This study |
| pAvA041 | X-4-p*PGK1*-*esaR*-t*ADH1* | This study |
| pAvA063 | XI-3-p*GAL1*-5x*luxO*-*amdS*-mKate2-t*CPS1* | [2] |
| pAvA070 | XI-1-t*ADH1*-At4CL1-p*ACT1*--p*PGI1*-FjTAL-tCYC | This study |
| pAvA079 | X-2-p*TDH3*-*esaI*_co-t*CYC1* | [2] |
| pAvA081 | X-2-p*TDH3*-*rpaI*_co-t*CYC1* | This study |
| pAvA103 | X-4-p*PGK1*-*GAL4*$_{AD}$-*luxR*-t*ADH1* | [2] |
| pAvA118 | XII-5-t*ADH1*-*SAM2*-p*TEF1*--p*PGK1*-*MET6*-t*CYC1* | [2] |
| pAvA122 | XI-1-t*ADH1*-At4CL1-p*ACT1*--p*PGI1*-AtPAL-t*CYC1* | This study |
| pAvA123 | XI-1-t*ADH1*-At4CL2-p*ACT1*--p*PGI1*-AtPAL-t*CYC1* | This study |
| pAvA124 | XI-1-t*ADH1*-At4CL2-p*ACT1*--p*PGI1*-FjTAL-t*CYC1* | This study |
| pAvA133 | XI-3-p*TEF1*_2x*esaO*_98_105-yeGFP-t*CPS1* | This study |
| pAvA139 | X-4-p*PGK1*-*GAL4*$_{AD}$-*luxR*_gen1 -t*ADH1* | [2] |
| pAvA145 | XI-3-p*GAL1*-5x*rpaO*-yeGFP-t*CPS1* | This study |
| pAvA146 | XII-5-p*GAL1*-5x*rpaO*-*amdS*-t*CPS1* | This study |
| pAvA155 | VIII-1-t*ADH1*-*SAM2*-p*TEF1*--p*PGK1*-*MET6*-t*CYC1* | This study |
| pAvA157 | X-4-p*PGK1*-*esaR*_D91G-NLS-t*ADH1* | This study |
| pAvA158 | X-2-p*TDH3*-*bjaI*_co-t*CYC1* | This study |
| pAvA159 | X-2-p*TDH3*-braI_co-t*CYC1* | This study |
| pAvA160 | XI-3-p*CCW*-*esaO*-yeGFP-t*CPS1* | This study |
| pAvA161 | XI-1-t*ADH1*-*SAM2*-p*TEF1*--p*PGK1*-*MET6*-t*CYC1* | This study |
| pAvA169 | X-4-p*PGK1*-VP48-NLS-*rpaR*-t*ADH1* | This study |
| pAvA171 | X-4-p*PGK1*-VP48-NLS-*traR*-t*ADH1* | This study |
| pAvA173 | X-4-p*PGK1*-VP48-NLS-*lasR*-t*ADH1* | This study |
| pAvA175 | X-2-p*TDH3*-*mesI*_co-t*ADH1* | This study |

| | | |
|---|---|---|
| pAvA176 | X-2-p*TDH3*-*mplI*_co-t*ADH1* | This study |
| pAvA179 | XI-3-p*GAL1*-5x*luxO*-mKate2-t*CPS1* | This study |
| pAvA185 | XI-3-p*GAL1*-5x*traO*-yeGFP-t*CPS1* | This study |
| pAvA187 | XI-3-p*GAL1*-5x*lasO*-yeGFP-t*CPS1* | This study |
| pAvA195 | XII-5-p*GAL1*-5x*traO*-*AmdS*-t*CPS1* | This study |
| pAvA197 | XII-5-p*GAL1*-5x*luxO*-mKate2-t*CPS1* | This study |
| pAvA199 | X-2-p*TDH3*-*mesI*-PRX-t*CPS1* | This study |
| pAvA200 | X-2-p*TDH3*-*bjaI*-PRX-t*CPS1* | This study |
| pAvA201 | X-2-p*TDH3*-braI-PRX-t*CPS1* | This study |
| pAvA202 | X-2-p*TDH3*-*rpaI*-PRX-t*CPS1* | This study |

**Table S4: overview of primers used in this study to create linear repair fragments for genomic integrations**

| Name | Target | Sequence |
|---|---|---|
| AA153 | *luxR*_N86K_fw | GAATGATTAGACTTGGAGTAATCAACGATTGGGTCG |
| AA154 | *luxR*_N86K_rv | CGTTGATTACTCCAAGTCTAATCATTCTCCCATCAACTGG |
| AA73 | p*PGK1*_ep_fw | GATCATCAAGGAAGTAATTATCTACTTTTTACAACAAATATAAAACAACCTGCACTAAAACAATG |
| AA74 | p*PGK1*_fw | GGTCTTTTCTAATTCGTAGTTTTTCAAGTTCTTAGATGCTTTCTTTTTCTCTTTTTTACAGATCATCAAGGAAGTAATTATCTACTTTTT |
| AA75 | p*PGK1*_rv | CATTGTTTTAGTGCAGGTTGTTTTATATTTG |
| AA76 | X-4_UP_fw | CCCAAAGCTAAGAGTCCCATT |
| AA77 | t*ADH1*_ep_rv | AATCATAAATCATAAGAAATTCGCTTATTTAGAAGTGTCAACAACGTATCTACATCTGTCCT |
| AA78 | t*ADH1*_rv | CTAAGAGTCACTTTAAAATTTGTATACACTTATTTTTTTTATAACTTATTTAATAATAAAAATCATAAATCATAAGAAATTCGCTTATTT |
| AA79 | t*ADH1*_fw | AGGACAGATGTAGATACGTTGTTG |
| AA80 | X-4_DN_rv | CTGGTGAGGATTTACGGTATGATC |

**Table S5: overview of primers used in this study for USER-cloning-based plasmids**

| name | target | sequence |
|---|---|---|
| MAD3 | BB_fw | ATCGCAUATCGCACGCATTCCATGCGA |
| MAD4 | BB_rv | ACCGCGTUATCGCGTACCAATTCGCCC |
| AA7 | *esaR*_fw | ACCTGCACUAAAACAATGTTCTCTTTCTTCCTTGAAAACCAAAC |
| AA8 | *esaR*_rv | ATCTGTCCUCTACCTTGCTGCTGACGCTG |
| AA256 | *esaR*-NLS_rv | ATCTGTCCUCTATACCTTTCTCTTCTTCTTTGGCCTTGCTGCTGACGCTG |
| AA291 | *lasR*_fw | AGAGAAAGGUATGGCCTTGGTTGACGGTTTTC |
| AA12 | *lasR*_rv | ATCTGTCCUTCAGAGAGTAATAAGACCCAAATTAACG |
| AA191 | *rpaR_fw* | AGAGAAAGGUATCGTCGGCGAAGATCAGC |
| AA21 | *rpaR*_rv | ATCTGTCCUTCACAAACGGATCAATCCGAGC |
| AA292 | *traR*_fw | AGAGAAAGGUATGCAGCACTGGCTGGAC |
| AA31 | *traR*_rv | ATCTGTCCUTTAGATCAGCTTTCTTCTGATTGCG |
| AA287 | VP48-NLS_fw | ACCTTTCTCUTCTTTGGGTGGACCTGGTAGCATATCAAGGTC |
| AA81 | *amdS*_fw | ACCTGCACUAAAACAATGCCACAATCTTGGGAAGAATTGG |
| AA121 | *amdS*_rv | ATCTGTCCUTTATGGAGTAACAACGTTACCCAAC |
| AA36 | p*GAL1*_core_fw | AGTGCGAUAAAACTGTATTATAAGTAAACTGAAAAAGCGTG |
| AA37 | p*GAL1*_core_rv | AGTGCAGGUGTCGACGCTAGCTATAGTTTTTTCTCC |
| AA245 | p*CCW*_fw | ACTTTATAUACTTGAACGGAGGCAAAGG |
| AA246 | p*CCW*-*esaO*_rv | ATATAAAGUCGGCAGCCTGTACTATAGTGCAGGTGGTGCTTGATAATCTTTCTTTCCATCC |
| AA247 | p*CCW*-*esaO*_fw | AGTGCAGGUTATTGATATAGTGTTTAAGCGAATGACAGAAG |
| AA248 | p*CCW*_rv | AGTGCGAUCGACACGCAAAAGAAAACCTTC |
| AA34 | p*TEF1*_fw | AGTGCGAUGCACACACCATAGCTTCAAAATG |
| AA35 | p*TEF1*_rv | AGTGCAGGUTTGTAATTAAAACTTAGATTAGATTGCTATGCTTTC |
| AA203 | p*TEF1*_2x*esaO*_rv | AAGACCGTUACCTGCACTATAGTACAGGCTATTAACTTAAATATCAATGGGAGGTCATCG |
| AA204 | p*TEF1*_2x*esaO*_fw | AACGGTCTUCGCCTGTACTATAGTG |

| | | |
|---|---|---|
| MAD14 | t*CPS1*_fw | AGGACAGAUGCGCAATGATTGAATAGTC |
| MAD15 | t*CPS1*_rv | AACGCGGUGTGGTTTTGATTTGATTTAAGTC |
| MAD36 | p*PGK1*_fw | AGTGCGAUAGACGCGAATTTTTCGAAGAAG |
| MAD37 | p*PGK1*_rv | AGTGCAGGUTGTTTTATATTTGTTGTAAAAAGTAGATAATTAC |
| MAD38 | p*TDH3*_fw | AGTGCGAUCTATTTTCGAGGACCTTGTCACC |
| MAD39 | p*TDH3*_rv | AGTGCAGGUTTTGTTTGTTTATGTGTGTTTATTCG |
| MAD61 | t*CYC1*_rv | AACGCGGUCTTCGAGCGTCCCAAAACCTTCTC |
| MAD62 | t*CYC1*_fw | AGGACAGAUATCCGCTCTAACCGAAAAGG |
| AA129 | p*ACT1*_fw | ATCGCACUGAAGCGGGTAAGCTGCCAC |
| AA130 | p*PGI1*_fw | AGTGCGAUAGAGAATTTTGCCATCGGACATGCTACCTTACGC |
| AA131 | *SAM2*_fw | AGTGCAGGUAAAACAATGTCCAAGAGCAAAACTTTCTTATTTAC |
| AA132 | *SAM2*_rv | CGTGCGAUTTAAAATTCCAATTTCTTTGGTTTTTCCC |
| AA144 | p*TEF1*_fw | ATCGCACUGCACACACCATAGCTTCAAAATG |
| AA145 | p*TEF1*_rv | ACCTGCACUTTGTAATTAAAACTTAGATTAGATTGCTATGCTTTC |
| AA161 | *MET6*_fw | ATCTGTCAUAAAACAATGGTTCAATCTGCTGTCTTAGGG |
| AA162 | *MET6*_rv | CACGCGAUTTAATTCTTGTATTGTTCACGGAAGTACTTG |
| AA96 | AtCL1 | AGTGCAGGUAAAACAATGGCTCCACAAGAACAAGCTG |
| AA97 | AtCL1 | CGTGCGAUTCACAAACCGTTAGCCAACTTGG |
| AA172 | At4CL2 | AGTGCAGGUAAAACAATGACTACCCAAGATGTTATCGTCAAC |
| AA173 | At4CL2 | CGTGCGAUTCAGTTCATCAAACCGTTAGCC |
| AA98 | FjTAL | ATCTGTCAUAAAACAATGAACACCATCAACGAATATCTGAGC |
| AA99 | FjTAL | CACGCGAUTTAATTGTTAATCAGGTGGTCTTTTACTTTCTGC |
| AA174 | AtPAL | ATCTGTCAUAAAACAATGGATCAAATCGAAGCTATGTTGTG |
| AA175 | AtPAL | CACGCGAUTCAGCAGATAGGAATAGGAGCAC |
| AA251 | *bjaI*_fw | ACCTGCACUAAAACAATGATCCATGCTATTTCTGCCG |
| AA252 | *bjaI*_rv | ATCTGTCCUTTAAGCAGACTTTCTTTGAGCAGC |
| AA253 | *braI*_fw | ACCTGCACUAAAACAATGCAAGCCACCATTAGAATCG |
| AA254 | *braI*_rv | ATCTGTCCUTTAAGCAGCAGCAGCCATTG |
| AA293 | *mplI*_fw | ACCTGCACUAAAACAATGATCACCGCTCATGTTGTTAACG |
| AA294 | *mplI*_rv | ATCTGTCCUTCATCTGGCAACTTCAGCCAAATG |
| AA295 | *mesI*_fw | ACCTGCACUAAAACAATGGTCAGAATCCATTTGGTTACCTG |
| AA296 | *mesI*_rv | ATCTGTCCUTCAAATGCCATCAGCTAAAATAGCACC |
| MAD12 | yeGFP_fw | ACCTGCACUAAAACAATGTCTAAAGGTGAAGAATTATTC |
| MAD13 | yeGFP_rv | ATCTGTCCUTTATTTGTACAATTCATCC |
| AA290 | mKate2_fw | ACCTGCACUAAAACAATGGTTTCTGAACTCATCAAGGAAAAC |
| AA86 | mKate2_rv | ATCTGTCCUTTATCTGTGTCCCAACTTAGATGGC |
| AA304 | *mesI*-PRX_rv | ATCTGTCCUTCACAATTTAGATCTCCTACCTCTTCCTAAAATGCCATCAGCTAAAATAGC ACC |
| AA306 | *rpaI*-PRX_rv | ATCTGTCCUTTACAATTTAGATCTCCTACCTCTTCCTAAAGAGATGACTTGGAATTCTGGC AAAG |
| AA307 | *bjaI*-PRX_rv | ATCTGTCCUTTACAATTTAGATCTCCTACCTCTTCCTAAAGCAGACTTTCTTTGAGCAGC |
| AA308 | *braI*-PRX_rv | ATCTGTCCUTTACAATTTAGATCTCCTACCTCTTCCTAAAGCAGCAGCAGCCATTG |

**Table S6: DNA sequences**

| Fragment | DNA sequence |
|---|---|
| VP48-NLS-*rpaR* | ATGCCAGCCGATGCTTTGGATGATTTCGATTTGGATATGTTGCCTGCTGATGCATTGGACG ATTTTGACTTAGACATGTTACCAGCAGACGCATTGGATGACTTCGACCTTGATATGCTACC AGGTCCACCcAAAGAAGAGAAAGGTATCGTCGGCGAAGATCAGCTTTGGGGACGGCGT GCGCTGGAATTCGTCGATTCCGTCGAACGGCTCGAGGCGCCGGCGCTGATCAGCCG GTTCGAATCGCTGATCGCGAGCTGCGGATTTACCGCCTACATCATGGCCGGCCTGCCG TCGCGCAATGCCGGACTACCGGAGCTGACGCTGGCCAATGGCTGGCCGCGAGACTG GTTCGATCTGTATGTCAGCGAAAACTTCAGCGCGGTCGATCCGGTGCCGCGCCACGGC GCTACCACGGTTCATCCTTTCGTATGGTCCGATGCACCCTACGACCGCGACCGTGATCC GGCCGCCCACCGGGTCATGACCCGGGCGGCGGAATTCGGACTGGTCGAGGGTTACT GCATTCCGCTGCACTACGACGACGGTAGCGCCGCGATCAGCATGGCCGGCAAGGATC CGGACCTCAGCCCGGCCGCGCGCGGCGCGATGCAGCTGGTCAGCATCTACGCGCA TAGTCGCCTGCGCGCACTCAGCCGGCCAAAGCCGATCCGGCGCAACCGGCTCACG |

| | |
|---|---|
| | CCGCGCGAGTGCGAGATCCTGCAATGGGCAGCGCAGGGCAAGACCGCCTGGGAAAT<br>CTCGGTAATCCTCTGCATCACCGAACGCACGGTGAAATTCCATCTGATCGAAGCCGCC<br>CGCAAGCTCGACGCCGCCAACCGCACCGCGGCGGTTGCCAAGGCATTGACGCTCG<br>GATTGATCCGTTTGTGA |
| VP48-NLS-*rpaR** | ATGCCAGCCGATGCTTTGGATGATTTCGATTTGGATATGTTGCCTGCTGATGCATTGGACG<br>ATTTTGACTTAGACATGTTACCAGCAGACGCATTGGATGACTTCGACCTTGATATGCTACC<br>AGGTCCACCcAAAGAAGAGAAAGGTATCGTCGGCGAAGATCAGCTTTGGGGACGGCGT<br>GCGCTGGAATTCGTCGATTCCGTCGAACGGCTCGAGGCGCCGGCGCTGATCAGCCG<br>GTTCGAATCGCTGATCGCGAGCTGCGGATTTACCGCCTACATCATGGCCGGCCTGCCG<br>TCGCGCAATGCCGGACTACCGGAGCTGACGCTGGCCAATGGCTGGCCGCGAGACTG<br>GTTCGATCTGTATGTCAGCGAAAACTTCAGCGCGGTCGATCCGGTGCCGCGCCACGGC<br>GCTACCACGGTTCATCCTTTCGTATGGTCCGATGCACCCTACGACCGCGACCGTGATCC<br>GGCCGCCCACCGGGTCATGACCCGGGCGGCGGAATTCGGACTGGTCGAGGGTTACT<br>GCATTCCGCTGCACTACGACGACGGTAGCGCCGCGATCAGCATGGCCGGCAAGGATC<br>CGGACCTCAGCCCGGCCGCGCGCGGCGCGATGCAGCTGGTCAGCATCTACGCGCA<br>TAGTCGCCTGCGCGCACTCAGCCGGCCAAAGCCGATCCGGCGCAACCGGCTCACG<br>CCGCGCGAGTGCGAGATCCTGCAATGGGCAGCGCAGGGCAAGACCGCCTGGGAAAT<br>CTCGGTAATCCTCTGCATCACCGAACGCACGGTGAAATTCCATCTGATCGAAGCCGCC<br>CGCAAGCTCGACGCCGCCAACCGCACCGCGGCGGTTGCCAAGGCATTGACGCTCG<br>GATTGATCCGTTTGTGA |
| VP48-NLS-*traR* | ATGCCAGCCGATGCTTTGGATGATTTCGATTTGGATATGTTGCCTGCTGATGCATTGGACG<br>ATTTTGACTTAGACATGTTACCAGCAGACGCATTGGATGACTTCGACCTTGATATGCTACC<br>AGGTCCACCcAAAGAAGAGAAAGGTATGCAGCACTGGCTGGACAAGTTGACCGATCTTG<br>CCGCAATTCAGGGCGACGAGTGCATCCTGAAGGATGGCCTTGCCGACCTTGCCGAAC<br>ATTTCGGCTTCACCGGCTATGCCTATCTCCATATCCAGCACAAACACACCATCGCGGTC<br>ACCAATTATCATCGTGACTGGCGATCGGCTTACTTCGAGAACAACTTCGACAAGCTCGAT<br>CCGGTCGTCAAGCGCGCGAAATCCAGGAAGCACGTCTTTGCCTGGTCCGGCGAACAG<br>GAACGATCGCGGCTATCGAAGGAAGAGCGTGCCTTCTACGCGCATGCGGCCGATTTCG<br>GCATCCGCTCCGGCATCACCATTCCGATCAAGACCGCCAACGGATCAATGTCGATGTT<br>CACGCTGGCGTCGGAAAGGCCGGCGATCGACCTCGACCGTGAGATCGACGCGGCC<br>GCAGCCGCGGGCGCCGTCGGGCAGCTCCATGCCCGCATCTCTTTCCTTCAGACCACT<br>CCGACAGTGGAAGATGCCGCCTGGCTCGATCCGAAAGAGGCGACCTATCTCAGATGGA<br>TCGCCGTCGGCATGACAATGGAGGAAGTCGCAGACGTGGAGGGCGTCAAGTACAACA<br>GCGTCCGTGTCAAGCTCCGCGAGGCCATGAAGCGCTTCGACGTTCGCAGCAAGGCC<br>CATCTCACCGCCCTCGCAATCAGAAGAAAGCTGATCTAA |
| VP48-NLS-*traR** | ATGCCAGCCGATGCTTTGGATGATTTCGATTTGGATATGTTGCCTGCTGATGCATTGaACG<br>ATTTTGACTTAGACATGTTACCAGCAGACGCATTGGATGACTTCGACCTTGATATGCTACC<br>AGGTCCACCcAAAGAAGAGAAAGGTATGCAGCACTGGCTGGACAAGTTGACCGATCTTG<br>CCGCAATTCAGGGCGACGAGTGCATCCTGAAGGATGGCCTTGCCGACCTTGCCGAAC<br>ATTaCGGCTTCACCGGCTtTGCCTATCTCCATATCCAGCACAAACACACCATCGCGGTCA<br>CCAATTATCATCGTGACTGGCGATCGGCTTACTTCGAGAACAACTTCGACAAGCTCGATC<br>CGGTCGTCAAGCGCGCGAAATCCAGGAAGCACGTCTTTGCCTGGTCCGGCGAACAGG<br>AtCGATCGCGGCTATCGAAGGAAGAGCGTGCCTTCTACGCGCgTGCGGCCGATTTCGG<br>CATCCGCTCCGGCATCACCATTCCGATCAAGACCGCCAACGGATCAATGTCGATGTTCA<br>CGCTGGCGTCGGAAAGGCCGGCGATCGACCTCGACCGTGAGATCGACGCGGCCGC<br>AGCCGCGGGCGCCGTCGGGCAGCTCCATGCCCGCATCTCTTTCCTTCAGACCACTCC<br>GACAGTGGAAGATGCCGCCTGGCTCGATCCGAAAGAGGCGACCTATCTCAGATGGATC<br>GCCGTCGGCATGACAATGGAGGAAGTCGCAGACGTGGAGGGCGTCAAGTACAACAGC<br>GTCCGTGTCAAGCTCCGCGAGGCCATGAAGCGCTTCGACGTTCGCAGCAAGGCCCAT<br>CTCACCGCCCTCGCAATCAGAAGAAAGCTGATCTAA |
| VP48-NLS-*lasR* | ATGCCAGCCGATGCTTTGGATGATTTCGATTTGGATATGTTGCCTGCTGATGCATTGGACG<br>ATTTTGACTTAGACATGTTACCAGCAGACGCATTGGATGACTTCGACCTTGATATGCTACC |

| | |
|---|---|
| | AGGTCCACCcAAAGAAGAGAAAGGTATGGCCTTGGTTGACGGTTTTCTTGAGCTGGAAC<br>GCTCAAGTGGAAAATTGGAGTGGAGCGCCATCCTGCAGAAGATGGCGAGCGACCTTGG<br>ATTCTCGAAGATCCTGTTCGGCCTGTTGCCTAAGGACAGCCAGGACTACGAGAACGCCT<br>TCATCGTCGGCAACTACCCGGCCGCCTGGCGCGAGCATTACGACCGGGCTGGCTAC<br>GCGCGGGTCGACCCGACGGTCAGTCACTGTACCCAGAGCGTACTGCCGATTTTCTGG<br>GAACCGTCCATCTACCAGACGCGAAAGCAGCACGAGTTCTTCGAGGAAGCCTCGGCC<br>GCCGGCCTGGTGTATGGGCTGACCATGCCGCTGCATGGTGCTCGCGGCGAACTCGGC<br>GCGCTGAGCCTCAGCGTGGAAGCGGAAAACCGGGCCGAGGCCAACCGTTTCATGGA<br>GTCGGTCCTGCCGACCCTGTGGATGCTCAAGGACTACGCACTGCAGAGCGGTGCCGG<br>ACTGGCCTTCGAACATCCGGTCAGCAAACCGGTGGTTCTGACCAGCCGGGAGAAGGA<br>AGTGTTGCAGTGGTGCGCCATCGGCAAGACCAGTTGGGAGATATCGGTTATCTGCAACT<br>GCTCGGAAGCCAATGTGAACTTCCATATGGGAAATATTCGGCGGAAGTTCGGTGTGACC<br>TCCCGCCGCGTAGCGGCCATTATGGCCGTTAATTTGGGTCTTATTACTCTCTGA |
| *bjaI* | ATGATCCATGCTATTTCTGCCGTTAACAGACACTTGTACGAAGATGTTTTGGAACAGCACTT<br>CAGATTGAGACACGATATCTTCGTTGAAGAAAGACATTGGGAAACTTTGAGAAGGCCAGA<br>TGGTAGAGAAGTTGATTCTTACGATGATGAAGATACCGTTTACTTGTTGGCTTTGGAAGGTA<br>GAAGAGTTGTTGGTGGTCATAGATTATACCCAACTACTAAGCCATCCATGATGTCTGAAGTT<br>TTTCCACATTTGGCTGCTGTTAGAGGTTGTCCATCTGATCCATTGATTTGGGAATGGTCTAG<br>ATACTTCGTTGTCAGGGATAGAAGAGATGGTGCTTTGAACTTGCAATTGATGGCTGCAGTT<br>CAAGAATTCTGTTTGGATCAAGGTATTGCTCAAGTTTCCGCTATTATGGAAACTTGGTGGTT<br>GCCAAGATTTCATGAAGCTGGTTTTGTTGTTACTCCATTGGGTTTGCCAGCTTTGGTTGAAA<br>ATGCTTGGACTATGGCTGCTACTGTTGATATTAGAAGGCAAACTTTGGATGTCTTGCACGAT<br>AGAATTGGTATGCCATCTATCGTTCAACAAGATGGTCCAAGATTGGATGCTGTTGCTAGAG<br>CTAATTTGTGTGGTTTAGCTGCTGCTCAAAGAAAGTCTGCTTAA |
| *braI* | ATGCAAGCCACCATTAGAATCGGTTTGAGACAAGAATTCGACAACGAACACATCAACGA<br>AATGTACAGATTGAGAGCCAGAGTTTTCAGAGATAGAATGGGTTGGGATATTCCAACTATT<br>GCCGGTATGGAAATTGATGGTTATGATGCTTTGGGTCCACACTACATGTTGATTCAAGAAC<br>CTGCTGGTGAAGTTAGAGGTTGTTGGAGATTGATGCCAACTGAAGGTCCAAATATGTTGAG<br>AGATACTTTCCCACAATTATTGGCTGGTAAAGCTGCTCCAACTGGTAGAACTATTTGGGAA<br>TTGTCTAGATTCGCTATTGAAGCTGGTGGTGATCAATCTTTTGGTTTCGCTGATGTTACCATG<br>CATGCTATTCATGCTTTGGTTACCTTTGCTGACAGAATGGGTATTACTAGATACGTTACTGTT<br>ACCACCACTCCAATTGAAAGGTTGTTGAGAAGAACCGGTATTGAATTAGCTAGATTAGGTC<br>CACCAATGAGAATTGGTACTGAAAACGCTATTGCTTTGGATATTGCTGTTTCTCCACAAAC<br>CAGAGTTGCTTTGTTTGGTCCAATGGCTGCTGCTGCTTAA |
| *mplI* | ATGATCACCGCTCATGTTGTTAACGCTCAAAACAGACACTTGTACGAAAACGAATTCGAC<br>GAATTTTTGAGAAGAAGGCACGATTTCTTCGTTCATCAAAAGAGATGGCGTCCACCATCTC<br>CAGATGGTAGAGAATTGGATCAATTTGATACTGATGCTGCCACCTACTTGTTAGGTATGGAA<br>GAAGGTAGAGTTGTTACCTCCGCTAGATTGATTCCAACTTCTGAACCACATATGGTGTCTG<br>AAGTTTTCCCTCATATGTGTGAAAAATCTGGTGTTCCAAGAAGGCCAGATTGGGCTGAATG<br>GACTAGAACTTTTGTTGTTCCAGAAAAGAGATCCACTGGTTTGAGAGGTACTTTGACTCAAT<br>TGTGTTGTGCCGTTATGGAATACGCTTTGGATGAAGGTTTGTCTGCTGTTGGTGGTGTACAA<br>GAAACTTACTTTATGCCACATCATGGTGCCTTGAAATGGCATGCTAAACCACTAGGTATGG<br>CTAGAGAAGAAAATGGTGAATGGTATATCGTTGCCTACATCGAAGTTAATGAAGCTGCTTTG<br>GCTTCCGTCAGAAAGATTTTGGGTTTAGACAGATCCTTGTTGGTCAGAAGAGGTATTCAATT<br>GCCATTCGTTAAGGTTAACAGACATTTGGCTGAAGTTGCCAGATGA |
| *mesI*-PRX | ATGGTCAGAATCCATTTGGTTACCTGGGAAAACAGAAAGTTGTACAGAAAGGTCTTGGAA<br>AGGTACTTCAGAATCAGATACGATATCTACGTCAAGCAAAGACGTTGGAGAGCTGTTGCTA<br>GACCTATTAACATTGAAATCGATGCCTTCGATAACGAACATGCCTTGTATGTTTTGGCTTTG<br>GATGCTAATGGTAAGATCGTTGGTGGTTCTAGATTGGTTCCAACATTAGAACCACACTTGAT<br>GTCTGAAGTTTTCCCAATTTTGGCTGGTGGTACTCCACCAAGAGCTGCTGAAATTTTTGAA<br>TGGACTAGATTCTTCGTCATGCCATCCTTTAGAACAAAAGGTGCTTCTTCACCAGTTGCTG<br>GTTTTGTTTTGTGTGGTTTGTTGGAAACTGCTCAGTCTTTGGGTATTAGACAAATCTCTGTTGT<br>TTGCGAAACTTTCTGGCCAAAAAGATTGAGAGCTTTAGGTTGGACTTTGTTCGAATTGGGT |

| | |
|---|---|
| | AATGCTTTGGAACATCCAGATGGTGATATTATCGCCTTGTTGATTGATGTTACCCCAGAAGC<br>TATTGAACAAACTAGACGTGCTTATGGTATTTCCGGTGCTATTTTAGCTGATGGCATTTTAGG<br>AAGAGGTAGGAGATCTAAATTGTGA |
| p*GAL*_5x*rpaO* | AAAACTGTATTATAAGTAAACTGAAAAAGCGTGTTTTTTATTCCTTAAGTTACGCAAGGTCGA<br>ACATAAGTACCTGTCCGATCGGACAGTATTACGCAAGGTCGAACATAAGTACCTGTCCGA<br>TCGGACAGTACGCTATTACGCCAGCGGATCCACCTGTCCGATCGGACAGTAGAACCTGT<br>CCGATCGGACAGTACAACCTGTCCGATCGGACAGTAACGTATGTGATCCGGATGATAATG<br>CGATTAGTTTTTTAGCCTTATTTCTGGGGTAATTAATCAGCGAAGCGATGATTTTTGATCTATT<br>AACAGATATATAAATGCAAAAACTGCATAACCACTTTAACTAATACTTTCAACATTTTCGGTT<br>TGTATTACTTCTTATTCAAATGTAATAAAAGTATCAACAAAAAATTGTTAATATACCTCTATACTT<br>TAACGTCAAGGAGAAAAAACTATAGCTAGCGTCGAC |
| pGAL_5x*lasO* | AAAACTGTATTATAAGTAAACTGAAAAAGCGTGTTTTTTATTCCTTAAGTTACGCAAGGTCGA<br>ACATAAGTATCTATCTCATTTGCTAGTTTTACGCAAGGTCGAACATAAGTATCTATCTCATTTG<br>CTAGTTCGCTATTACGCCAGCGGATCCATCTATCTCATTTGCTAGTTGAATCTATCTCATTTG<br>CTAGTTCAATCTATCTCATTTGCTAGTTACGTATGTGATCCGGATGATAATGCGATTAGTTTTT<br>TAGCCTTATTTCTGGGGTAATTAATCAGCGAAGCGATGATTTTTGATCTATTAACAGATATATA<br>AATGCAAAAACTGCATAACCACTTTAACTAATACTTTCAACATTTTCGGTTTGTATTACTTCTT<br>ATTCAAATGTAATAAAAGTATCAACAAAAAATTGTTAATATACCTCTATACTTTAACGTCAAGG<br>AGAAAAAACTATAGCTAGCGTCGAC |
| pGAL_5x*traO* | AAAACTGTATTATAAGTAAACTGAAAAAGCGTGTTTTTTATTCCTTAAGTTACGCAAGGTCGA<br>ACATAAGTATGTGCAGATCTGCACATTTACGCAAGGTCGAACATAAGTATGTGCAGATCTG<br>CACATCGCTATTACGCCAGCGGATCCATGTGCAGATCTGCACATGAATGTGCAGATCTGC<br>ACATCAATGTGCAGATCTGCACATACGTATGTGATCCGGATGATAATGCGATTAGTTTTTTA<br>GCCTTATTTCTGGGGTAATTAATCAGCGAAGCGATGATTTTTGATCTATTAACAGATATATAA<br>ATGCAAAAACTGCATAACCACTTTAACTAATACTTTCAACATTTTCGGTTTGTATTACTTCTTA<br>TTCAAATGTAATAAAAGTATCAACAAAAAATTGTTAATATACCTCTATACTTTAACGTCAAGGA<br>GAAAAAACTATAGCTAGCGTCGAC |
| p*CCW_esaO* | CGACACGCAAAAGAAAACCTTCGAGGTTGCGCACTTCGCCCACCCATGAACCACACG<br>GTTAGTCCAAAAGGGGCAGTTCAGATTCCAGATGCGGGAATTAGCTTGCTGCCACCCTC<br>ACCTCACTAACGCTGCGGTGTGCGGATACTTCATGCTATTTATAGACGCGCGTGTCGGAA<br>TCAGCACGCGCAAGAACCAAATGGGAAAATCGGAATGGGTCCAGAACTGCTTTGAGTG<br>CTGGCTATTGGCGTCTGATTTCCGTTTTGGGAATCCTTTGCCGCGCGCCCCTCTCAAAA<br>CTCCGCACAAGTCCCAGAAAGCGGGAAAGAAATAAAACGCCACCAAAAAAAAAAATAAA<br>AGCCAATCCTCGAAGCGTGGGTGGTAGGCCCTGGATTATCCCGTACAAGTATTTCTCAG<br>GAGTAAAAAAACCGTTTGTTTTGGAATTCCCCATTTCGCGGCCACCTACGCCGCTATCTT<br>TGCAACAACTATCTGCGATAACTCAGCAAATTTTGCATATTCGTGTTGCAGTATTGCGATAA<br>TGGGAGTCTTACTTCCAACATAACGGCAGAAAGAAATGTGAGAAAATTTTGCATCCTTTGC<br>CTCCGTTCAAGTATATAAAGTCGGCAGCCTGTACTATAGTGCAGGTGGTGCTTGATAATCTT<br>TCTTTCCATCCTACATTGTTCTAATTATTCTTATTCTCCTTTATTCTTTCCTAACATACCAAGAA<br>ATTAATCTTCTGTCATTCGCTTAAACACTATATCAATAACCTGCACT |
| *esaR*$_{D91G}$-NLS | TTCTCTTTCTTCCTTGAAAACCAAACAATAACGGATACGCTTCAGACTTACATACAGAGAAA<br>GTTATCTCCGCTGGGTAGTCCGGATTACGCTTACACTGTTGTGAGCAAAAAAAATCCTTCA<br>AATGTTCTGATTATTTCCAGTTATCCTGACGAATGGATTAGGTTATACCGCGCTAACAACTTT<br>CAGCTGACCGATCCGGTTATTCTCACGGCCTTTAAACGCACCTCGCCGTTTGCCTGGGA<br>TGAGAATATTACGCTGATGTCCGACCTGCGGTTCACCAAAATTTTCTCTTTATCCAAGCAAT<br>ACAACATCGTTAACGGCTTTACCTATGTCCTGCATGACCACATGAACAACCTTGCTCTGTT<br>GTCCGTGATCATTAAAGGCAACGATCAGACTGCGCTGGAGCAACGCCTTGCTGCCGAA<br>CAGGGCACGATGCAGATGCTGCTGATTGATTTTAACGAGCAGATGTACCGCCTGGCAGG<br>CACCGAAGGTGAACGAGCCCCGGCGTTAAATCAGAGCGCGGACAAAACGATATTTTCC<br>TCGCGTGAAAATGAGGTGTTGTACTGGGCGAGTATGGGCAAAACCTATGCTGAGATTGCC<br>GCTATTACGGGCATTTCTGTGAGTACCGTGAAGTTTCACATCAAGAATGTGGTCGTGAAAC |

| | |
|---|---|
| | TGGGCGTCAGTAACGCCCGACAGGCTATCAGACTGGGTGTAGAACTGGATCTTATCAGACCGGCAGCGTCAGCAGCAAGGCCAAAGAAGAAGAGAAAGGTATAG |
| p*TEF1_esaO* | GCACACACCATAGCTTCAAAATGTTTCTACTCCTTTTTTACTCTTCCAGATTTTCTCGGACTCCGCGCATCGCCGTACCACTTCAAAACACCCAAGCACAGCATACTAAATTTCCCCTCTTTCTTCCTCTAGGGTGTCGTTAATTACCCGTACTAAAGGTTTGGAAAAGAAAAAAGAGACCGCCTCGTTTCTTTTTCTTCGTCGAAAAAGGCAATAAAAATTTTTATCACGTTTCTTTTTCTTGAAAATTTTTTTTTTTGATTTTTTTCTCTTTCGATGACCTCCCATTGATATTTAAGTTAATAAACGGTCTTCGCCTGTACTATAGTGCAGGTAATTTCTCAAGTTTCAGTTTCATTTTTCTTGTTCTATTACAACTTTTTTTACTTCTTGCTCATTAGAAAGAAAGCATAGCAATCTAATCTAAGTTTTAATTACAA |
| p*TEF1*_2x*esaO* | GCACACACCATAGCTTCAAAATGTTTCTACTCCTTTTTTACTCTTCCAGATTTTCTCGGACTCCGCGCATCGCCGTACCACTTCAAAACACCCAAGCACAGCATACTAAATTTCCCCTCTTTCTTCCTCTAGGGTGTCGTTAATTACCCGTACTAAAGGTTTGGAAAAGAAAAAAGAGACCGCCTCGTTTCTTTTTCTTCGTCGAAAAAGGCAATAAAAATTTTTATCACGTTTCTTTTTCTTGAAAATTTTTTTTTTTGATTTTTTTCTCTTTCGATGACCTCCCATTGATATTTAAGTTAATAGCCTGTACTATAGTGCAGGTAACGGTCTTCGCCTGTACTATAGTGCAGGTAATTTCTCAAGTTTCAGTTTCATTTTTCTTGTTCTATTACAACTTTTTTTACTTCTTGCTCATTAGAAAGAAAGCATAGCAATCTAATCTAAGTTTTAATTACAA |
| *esaI* | ATGTTGGAGTTGTTCGATGTCTCCTACGAAGAATTGCAAACTACTAGATCTGAAGAGTTGTACAAGCTGAGGAAAAAGACCTTCTCTGATAGATTAGGTTGGGAAGTTATTTGCTCCCAAGGTATGGAATCTGATGAATTTGATGGTCCAGGTACTAGGTACATTTTGGGTATTTGTGAAGGTCAATTGGTTTGCTCTGTTAGGTTCACTTCTTTGGATAGACCAAACATGATTACCCATACCTTCCAACACTGTTTCTCTGATGTTACTTTGCCAGCTTATGGCACTGAATCTTCTAGATTCTTTGTTGATAAGGCTAGAGCTAGAGCTTTGTTGGGTGAACATTATCCAATCTCTCAAGTTTTGTTCTTGGCCATGGTTAATTGGGCTCAAAACAATGCTTACGGTAACATCTACACCATCGTTTCAAGAGCCATGTTGAAGATTTTGACAAGATCTGGTTGGCAGATCAAGGTTATCAAAGAAGCTTTCTTGACCGAGAAAGAGAGGATCTATTTGTTGACATTGCCAGCTGGTCAAGATGACAAACAACAATTAGGTGGTGATGTTGTTTCTAGAACTGGTTGTCCACCAGTTGCTGTTACTACTTGGCCATTGACTTTACCAGTTTAA |
| *rpaI* | ATGCAAGTTCACGTTATCAGAAGAGAAAACAGAGCATTATACGCTGGTTTGCTGGAAAAGTACTTCAGAATCAGACACCAAATCTACGTTGTTGAGAGAGGTTGGAAAGAATTGGATAGACCAGATGGTAGAGAAATCGATCAATTCGATACTGAAGATGCCGTTTACTTGTTGGGTGTTGATAACGATGATATCGTTGCTGGTATGAGAATGGTTCCAACTACTTCTCCAACTTTGTTGTCTGATGTTTTCCCACAATTGGCTTTGGCTGGTCCAGTTAGAAGGCCAGATGCTTATGAATTGTCCAGAATTTTCGTTGTCCCAAGAAAAAGGGGTGAACATGGTGGTCCAAGAGCTGAAGCTGTTATTCAAGCTGCTGCTATGGAATACGGTTTGTCTATTGGTTTGTCTGCTTTCACCATCGTTTTGGAAACTTGGTGGTTGCCAAGATTGGTTGATCAAGGTTGGAAGGCTAAACCATTGGGTTTGCCACAAGATATTAACGGTTTTTCTACTACCGCCGTTATCGTTGATGTTGATGATGATGCTTGGGTTGGTATCTGTAACAGAAGATCTGTTCCAGGTCCAACTTTAGAATGGCGTGGTTTGGAAGCTATTAGAAGGCATTCTTTGCCAGAATTCCAAGTCATCTCTTAA |
